# Spatiotemporal properties of cortical excitatory and inhibitory neuron activation by sustained and bursting electrical microstimulation

**DOI:** 10.1101/2024.09.30.615029

**Authors:** Christopher L Hughes, Kevin A Stieger, Keying Chen, Alberto L Vazquez, Takashi DY Kozai

**Author notes:** denotes equal contributions (Christopher L Hughes, Kevin A Stieger, Keying Chen).

## Abstract

Intracortical microstimulation (ICMS) of sensory cortices produces artificial sensation yet the neural mechanisms underlying evoked responses, particularly among inhibitory subpopulations, remain unclear. We investigated how long durations (30 s) of ICMS shape spatiotemporal patterns in excitatory and inhibitory network activation using two-photon imaging of visual cortex in transgenic mice. Inhibitory neurons were more likely to facilitate (increase in activation) across 30 s of ICMS, whereas excitatory neurons were more likely to adapt (decrease in activation) and exhibit post-ICMS depression. Different temporal profiles led to preferential activation of excitatory or inhibitory neurons, with theta-burst stimulation driving the strongest inhibitory response and 10-Hz burst patterns driving the strongest peak excitatory response. Neurons located farther from the electrode exhibited more diverse responses to ICMS, highlighting synaptic recruitment dynamics such as inhibition and disinhibition. This study reveals how ICMS differentially influences excitatory and inhibitory neuron activity across long durations of ICMS and suggests temporal patterning can be used to potentially target neuronal subpopulations and drive desirable activity patterns.

## Introduction

Intracortical microstimulation (ICMS) focally activates neurons within the cerebral cortex and has recently been applied for artificial sensory restoration in clinical research^1–5^. Emerging neurotech companies promise to ameliorate neurological conditions and provide autonomy to affected clinical populations by recording and stimulating on implanted electrodes^6–9^. However, much remains unclear about how ICMS affects the brain. In recent studies, ICMS in visual cortex restored rudimentary vision in a person^5^ and, with higher electrode density, produced discriminable shapes in the visual field of non-human primates^10^. Nevertheless, vision restored with ICMS is still far less functional than normal vision. Understanding ICMS mechanisms will be crucial for improving therapeutic outcomes.

Studying the neural mechanisms of ICMS is difficult in humans due to hardware limitations and challenges to recording neural activity during stimulation in addition to limitations in spatial resolution. Recently, two-photon microscopy in transgenic mouse models provided unprecedented insight into how ICMS drives neuronal activity^11–17^. In these models, higher frequencies of ICMS adapted activity far from the electrode, but lower frequencies were able to continue to recruit neurons far away from the electrode over time^12–15^. Additionally, individual neurons responded preferentially to different temporal profiles or waveform shapes, with some neurons responding more to low frequencies, high frequencies, or burst profiles^14,15^. Transgenic mouse models can then contribute to understanding how ICMS parameters affect neural recruitment and inform therapeutic approaches.

The role of inhibitory neurons in ICMS recruitment remains underexplored^17^. Inhibition is fundamental to cortical sensory processing and plays important roles in encoding sensory specific features, such as contrast enhancement and increased sensitivity to moving objects^18–26^. One recent study investigated inhibitory neuron responses to ICMS^17^, but this study focused on short 100 ms durations that are not representative of clinical sensory restoration applications that require long or continuous stimulation. Most clinical research to date in somatosensory restoration has focused on train durations of at least 1 s and up to 60 s^2,27–30^. For intracortical visual prosthesis, work has focused on durations of 200 ms, which provided perception of brief phosphenes but did not allow for continuous visual perception^5^. Unfortunately, longer durations (>20 s in S1 and a few seconds in V1) of ICMS consistently drive adaptation in a stimulation amplitude and frequency dependent manner for both evoked percepts in humans as well as evoked neural activity in mice^12–15,31,32^.

One hypothesis is that inhibitory neurons contribute to the adaptation of neural activity, but investigating the contribution of inhibitory neurons to induced adaptation requires longer stimulation trains. Additionally, increases in blood supply to the ICMS activated region results in elevated oxyhemoglobin after ICMS-activity has stabilized, correlating with prolonged neuronal adaptation (10-20 s after ICMS onset)^32^. Inhibitory neurons play a strong role in regulation of the blood flow^33^, further suggesting the importance of studying longer ICMS trains. Additionally, GCaMP measured activity has long latencies between neural activation and signal detection (300-500 ms^34,35^), so that longer trains provide more insight about the overall activation and excitatory-inhibitory dynamics that can be achieved with continuous ICMS. These reasons motivated examination of the response of excitatory and inhibitory populations to longer, more clinically-relevant ICMS durations without the influence of anesthesia. Here, we investigated how excitatory and inhibitory neuronal subtypes respond to long ICMS trains under awake conditions.

## Results

### Investigating the role of inhibitory neurons in response to long ICMS trains

In ICMS applications for sensory restoration, low-frequency stimulation evokes continuous tapping sensations, while high-frequency stimulation leads to rapid sensory decline within the first few seconds^2,28^. Inhibitory neurons are crucial in sensory processing, and likely play a role in these ICMS-evoked perceptual responses^19,25,36–39^. To enhance clinical therapies, it is essential to determine how inhibitory neurons influence longer ICMS-evoked responses. Here, we explore how different temporal profiles affect excitatory and inhibitory neuron activation. Our previous observations showed significant changes in the spatiotemporal activation of ICMS-evoked excitatory network activity within the initial 10-20 s^11–15,32^. However, its impact on inhibitory neurons remains unclear. Given the prevalent use of prolonged ICMS in clinical settings, understanding the mechanisms driving network dynamics will be pivotal for improving ICMS-based clinical approaches.

To study how ICMS impacts excitatory and inhibitory neurons over longer, clinically relevant ICMS durations, mice expressing red fluorescence in inhibitory neurons (tdTomato) and green fluorescence for calcium evoked activity (GCaMP7b) in all neurons were implanted with Michigan probes in the visual cortex (Fig. 1A). We quantified GCaMP responses while stimulating one electrode in Layer 2/3 with four 30-s ICMS trains with different temporal patterns reflective of clinically applied trains (10 Hz uniform, 100 Hz uniform, 10 Hz Burst, and theta-burst stimulation (TBS); Fig. 1B-D). However, the fluorescence in the green channel contributed to fluorescence in the red channel (SFig. 1A). To separate GCaMP labeled neurons into excitatory and inhibitory subpopulations, we corrected for cross-channel fluorescence using linear regression (SFig. 1B). Neurons with high red intensity (neurons with a value greater than the model plus the root-mean squared error) were classified as “inhibitory,” those with low intensity (below the model line) were classified as “excitatory,” and neurons in between were classified as “uncertain.” After classification, 54% of neurons (333 total) were excitatory, 16% were inhibitory (68 total), and 30% were uncertain (158 total) (Fig. 2A). In cortex, typically ∼80% of neurons are excitatory while ∼20% of neurons are inhibitory, indicating that most of the neurons in the “uncertain” population represent excitatory neurons^40–43^. Throughout the paper, we refer to these different classifications as excitatory, inhibitory, and uncertain “subpopulations.”

**Figure 1:**
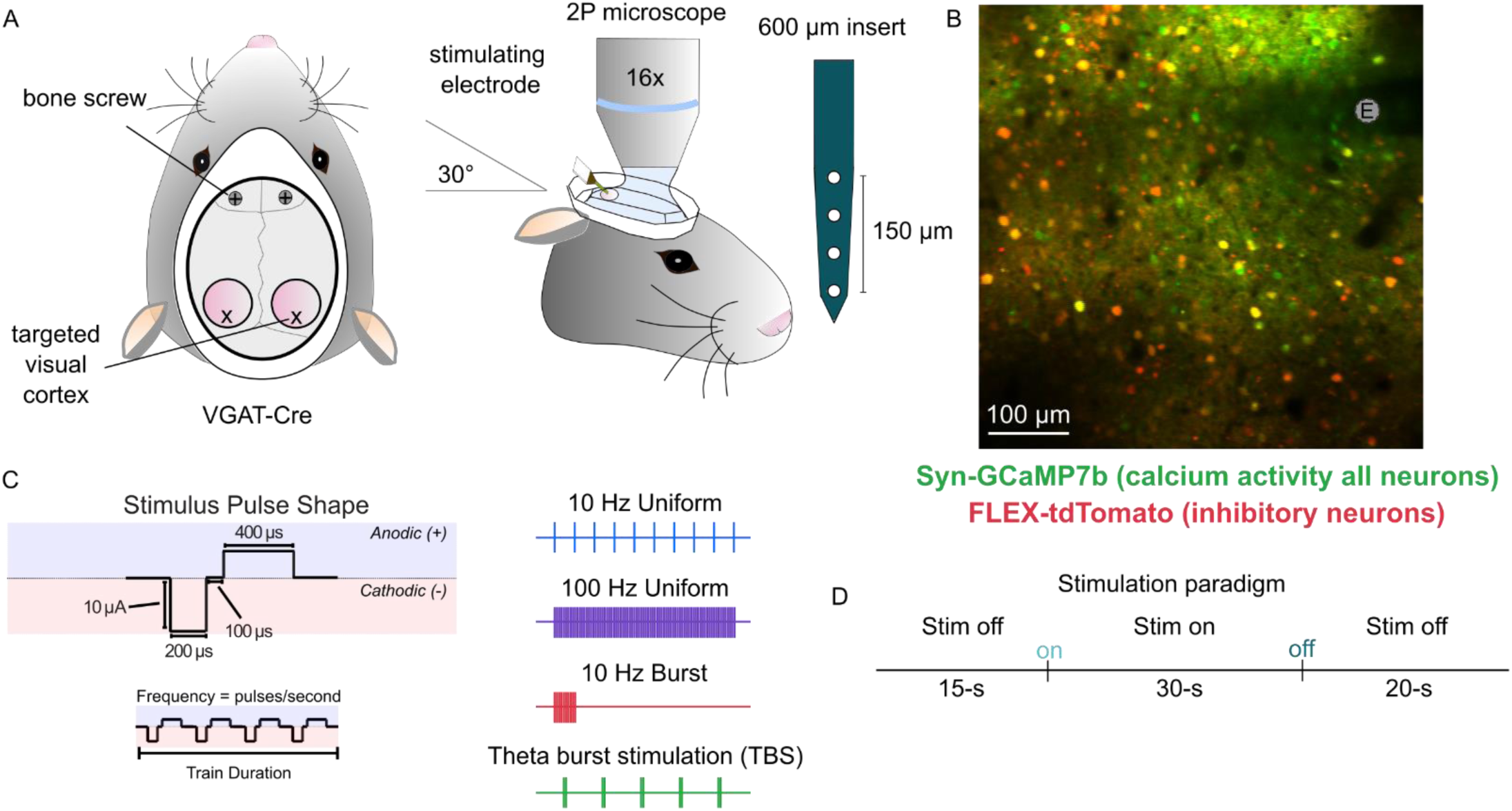
Two-photon imaging of VGAT-Ires-Cre mouse visual cortex in response to microstimulation. A) Bilateral craniotomies were made over visual cortex of VGAT-Cre mice, then AAV-FLEX-tdTomato and AAV-Syn-GCaMP7b were injected. After >3 weeks for expression, a Michigan probe was inserted into an area of expression while the animal was under light isoflurane anesthesia. All data collection occurred after the animal awoke. B) Example image of VGAT-expressing neurons (red) and calcium activity from all neurons (green) around the microelectrode (E). C) ICMS pulse shape and parameters. Four different ICMS trains were provided with different temporal profiles. D) Paradigm for ICMS delivery with 15 s pre-ICMS, 30 s ICMS, and 20 s post-ICMS imaging.

**Figure 2:**
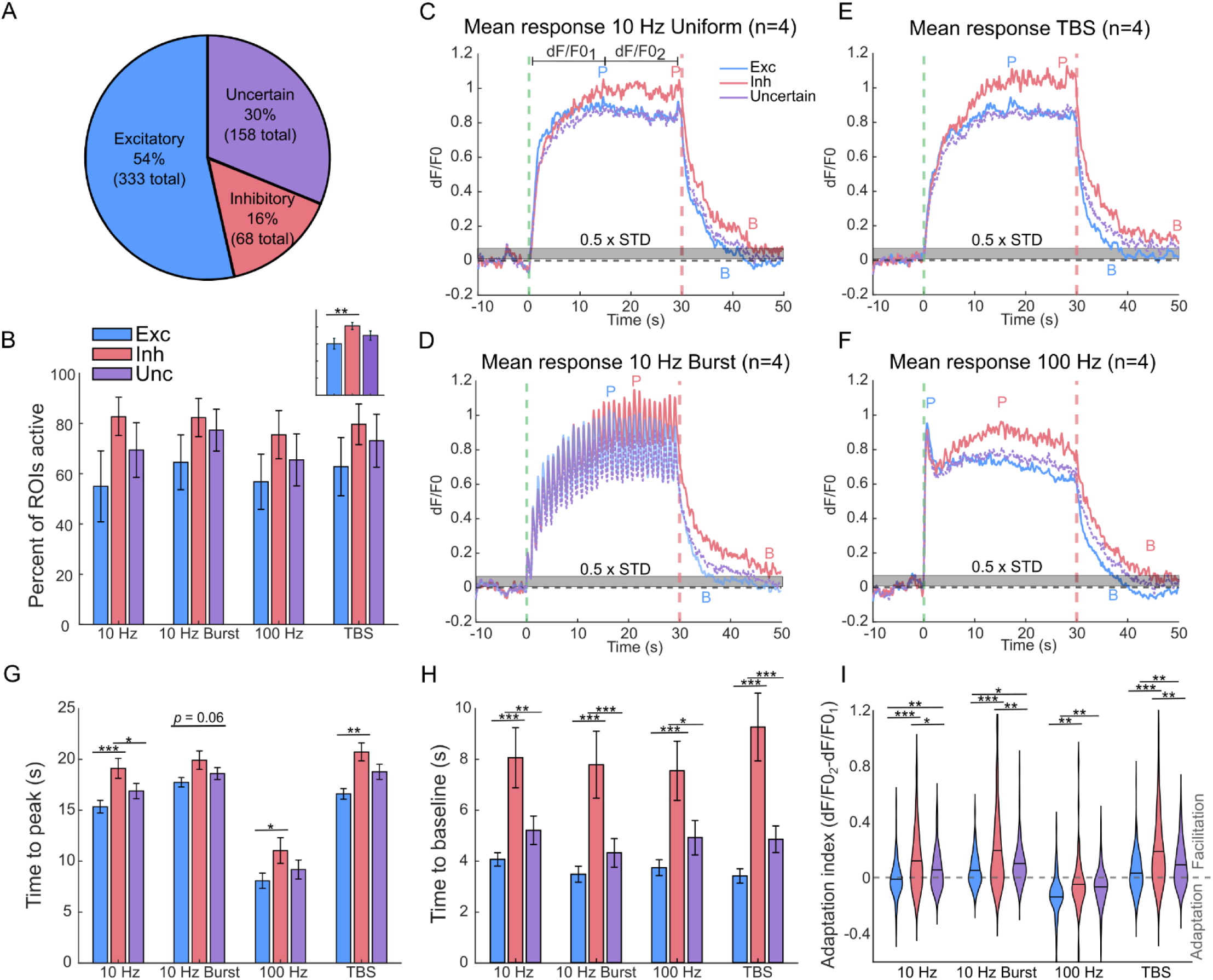
Inhibitory neurons facilitate more to ICMS and have slower time-to-peak and return-to-baseline than excitatory or uncertain neurons. A) The distribution of neurons labeled as excitatory (333 total), inhibitory (68 total), and uncertain (158 total). B) Total percent of neurons active during ICMS for each subpopulation across trains. Error bars represent standard error across animals (n=4). The inset shows the comparison of subpopulations combined across trains, with a higher percent of inhibitory neurons active. C-F) Mean traces of each subpopulation to 10 Hz Uniform (C), 10-Hz Burst (D), TBS (E), or 100 Hz (F) ICMS. ICMS start (green vertical line), ICMS stop (red vertical line), 0.5x the standard deviation of baseline activity (grey bar), detected time-to-peak (“P”), and detected time-to-baseline (“B”) are indicated for each subpopulation. The two intervals of dF/F01 and dF/F02 represent the intervals used for calculation of the adaptation in (I). G-H) The calculated time-to-peak (G) and time-to-baseline (H) for each subpopulation shows generally slower times for inhibitory neurons. I) The adaptation index for each subpopulation shows less adaptation/more facilitation for inhibitory neurons.

Neurons were also classified by their response to ICMS. “Active” neurons were defined as having activity greater than 3 standard deviations above baseline during ICMS and were otherwise classified as “inactive.” Among active neurons, those continuously active for more than 0.5 s were categorized as "consecutive active," while others were termed "non-consecutive active." This classification enabled the study of neurons displaying brief bouts of activity during ICMS. For simplicity, we refer to consecutive active as “active” and non-consecutive active as “non-consecutive” in the results. Overall, neurons tended to maintain their classification across stimulation parameters (SFig. 2). Transitions between classifications generally involved only one category change (between active and non-consecutive, or non-consecutive and inactive). For neurons that changed classes across trains, 10 Hz was more likely to drive inactivity or non-consecutive activity while TBS was more likely to drive consecutive activation. Inhibitory neurons demonstrated greater stability in activation across trains (SFig. 2).

### Inhibitory neurons were more strongly activated and facilitated by ICMS than excitatory neurons

Given the observed dynamics in network activation over extended stimulation periods^11–15,32^, we aimed to determine how longer-duration ICMS affects excitatory and inhibitory neurons. To understand the activation intensity for these subpopulations, the percentage of consecutively active neurons was evaluated in each subpopulation (Fig. 2B). Across the four ICMS trains, no significant differences were found in the percentage of activated neurons within any subpopulation (*p*>0.05). Inhibitory neurons consistently exhibited the highest percentage of active neurons (83±4%), whereas excitatory neurons showed the lowest (61±7%). This difference was statistically significant when aggregating data from all ICMS trains (*p*<0.01, Fig. 2B inset). These results indicate that while each ICMS train evoked a comparable number of active neurons overall, inhibitory neurons generally displayed higher population recruitment compared to excitatory neurons.

Temporal patterning of stimulation has been proposed as a method to modulate the strength and duration of activation of different recruited neural elements^15,44^. Given that percent activations were similar across trains, active neurons were evaluated for differences in the mean intensity evoked by each of the four ICMS trains. All low-frequency trains (10 Hz Uniform, 10-Hz Burst, and TBS) generally evoked activity that was stable or increasing/facilitating across the ICMS duration (Fig. 2C-E). 10-Hz burst trains also produced clear modulation with each burst (Fig. 2D). In contrast, 100-Hz trains evoked intense and rapidly adapting activation in the first 3 s of ICMS with more stable activation from 4-30 s (Fig. 2F). Across all trains, inhibitory neurons exhibited a more facilitated average response than excitatory neurons (Fig. 2C-F). Within the first 10-15 s, mean intensities overlapped for all subpopulations (Exc, Inh, Unc; *p*>0.05). However, beyond this period, inhibitory neurons consistently showed higher average intensities than excitatory and uncertain neurons. This difference in mean intensity was statistically significant only for the TBS train (*p*=0.036), which exhibited the largest disparity in mean intensity between inhibitory (mean dF/F0=1.05±0.08), excitatory (mean dF/F0=0.82±0.05), and uncertain (mean dF/F0=0.85±0.05) neurons. Different ICMS temporal patterns resulted in varying temporal responses across all subpopulations, with inhibitory neurons consistently demonstrating greater facilitation across all temporal patterns.

In both clinical applications and basic science research, long sequences of ICMS trains are continuously delivered. However, it remains unclear how ICMS parameters affect excitatory and inhibitory neuron recovery. To evaluate temporal progression in different subpopulations, significant changes in ‘time-to-peak’, ‘return-to-baseline’, and ‘overall change’ in activity across the train was evaluated. Inhibitory neurons exhibited prolonged facilitation later into the trains, leading to later peaks and higher activity levels at the end of the train. Consequently, inhibitory neurons showed slower time-to-peak (Fig. 2G) and slower return-to-baseline (Fig. 2H) than excitatory neurons. The 100-Hz train induced a faster time-to-peak across all subtypes due to rapid adaptation (*p*<0.001), but no difference in post-ICMS return-to-baseline (*p*>0.05). Across all trains, inhibitory neurons ‘facilitated more’/’adapted less’ in fluorescent intensity than excitatory neurons as indicated by the adaptation index (Fig. 2I). Moreover, the 100-Hz trains evoked significantly ‘less facilitation’/’more adaptation’ in evoked fluorescence intensity across all subpopulations (*p*<0.001). For excitatory neurons, 10-Hz burst trains evoked significantly more facilitation than 10-Hz uniform trains (p<0.001); however, lower activity was also observed compared to other low-frequency stimulation due to inhibition during the off-period of the burst cycle. Together these results indicate that high frequencies lead to faster time-to-peak, but all trains exhibit similar return-to-baseline. Inhibitory neurons had more facilitation than excitatory neurons resulting in stronger activity throughout 30-s ICMS for all trains, with the burst paradigms evoking the strongest facilitation overall. Notably, differences between excitatory and inhibitory neurons occurred after 10-15 s of ICMS. This underscores the importance of longer stimulation durations on inhibitory neurons, which are undetectable with shorter train durations used in previous work^17^. Low-frequency trains evoked stable or facilitating responses in all subpopulations, while the 100-Hz train induced robust initial activation followed by pronounced adaptation, consistent with previous findings^12–15^. Here, we show inhibitory neurons strengthen activity over time to prolonged ICMS, which can impact network activity by adapting excitatory neurons^12–15,32,45^.

### Individual excitatory and inhibitory neurons have preferential activation by different temporal patterns

We showed that inhibitory neurons were more likely to facilitate and have higher evoked intensity than excitatory neurons for all presented ICMS trains. However, it remains unclear whether any of the four ICMS trains elicited stronger responses in excitatory or inhibitory neurons, and if individual neurons exhibited differential responses to each train. To investigate these dynamics and their implications for altering network activation patterns, we sought to determine the preferential responses of individual excitatory and inhibitory to the four presented trains by assessing the evoked fluorescent intensity (Fig. 3A).

**Figure 3:**
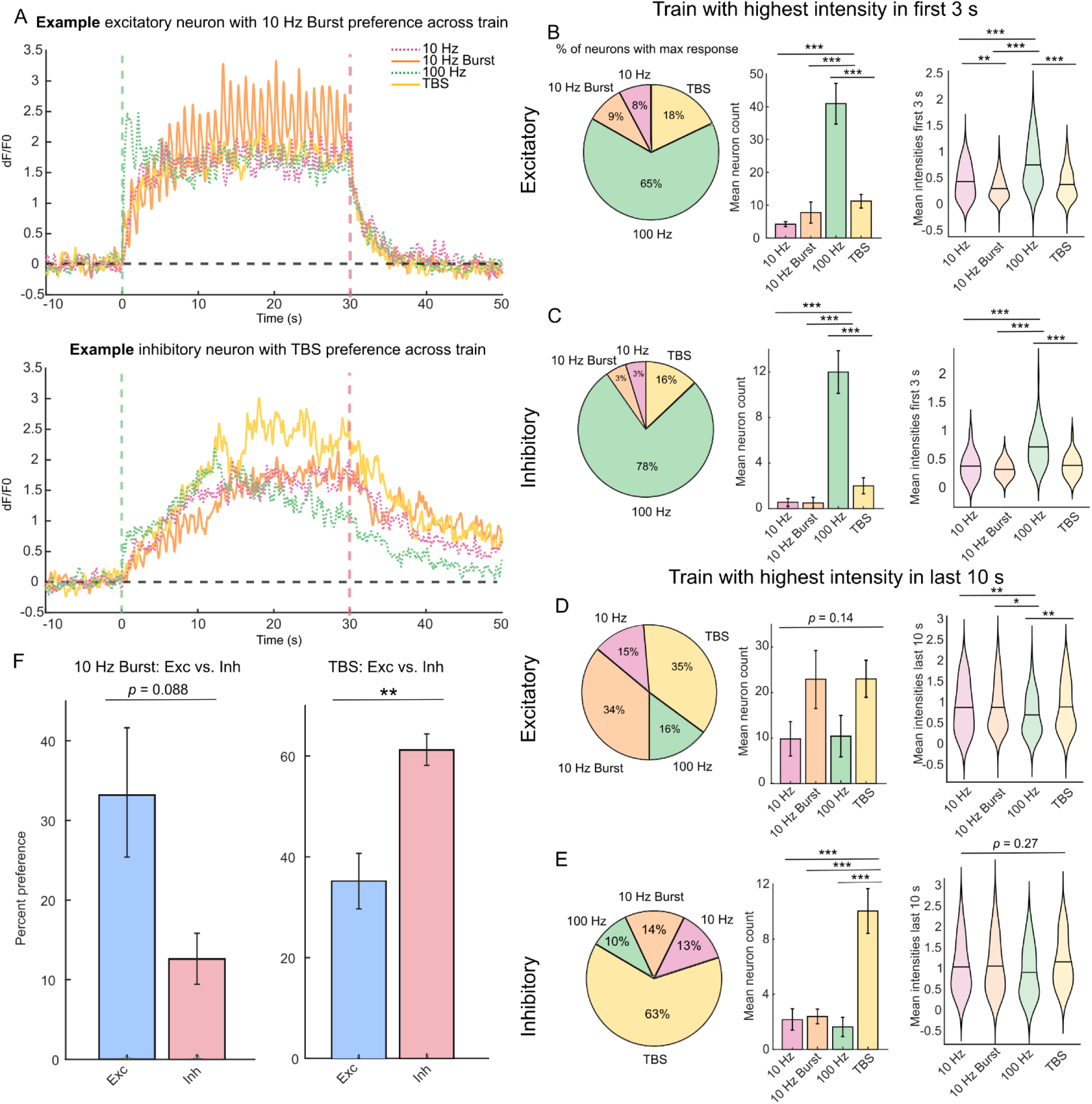
TBS trains drive stronger activation of individual inhibitory neurons, while 10-Hz Burst trains drive stronger activation of individual excitatory neurons. A) Examples of an individual excitatory neuron with preference to 10-Hz Burst (top) and an individual inhibitory neuron with preference to TBS (bottom). Vertical bars represent ICMS start (green) and stop (red). B-C) Preferential responses of excitatory (B) and inhibitory (C) neurons in the first 3 s of ICMS. Most neurons had the strongest response to 100 Hz. Pie plot shows the percent of each subpopulation with the highest intensity for each train. Bar plots show the total neuron counts with the highest intensity for each train. Error bars indicate standard error across animals. Violin plots show the distribution of mean intensities across all active neurons. D-E) Preferential responses of excitatory (D) and inhibitory (E) neurons in the last 10 s of ICMS. Excitatory neurons show strong preferences for 10-Hz Burst and TBS, whereas inhibitory neurons predominantly preferred TBS. F) Bar plots indicating preference for 10-Hz Burst trains (left) and TBS trains (right) in the last 10 s of ICMS for excitatory vs. inhibitory neurons. Error bars represent the standard error across animals.

In the previous section and studies^12–15^, ICMS responses in the initial few seconds differ from those in the subsequent period, suggesting a shift in excitatory-inhibitory activation over time. Therefore, we analyzed preferential responses during two distinct time windows: the first 3 s (Fig. 3B-C) and the last 10 s after stabilization (Fig. 3D-E). During the first 3 s, most neurons across all subpopulations exhibited the highest intensity response to 100-Hz trains (*p*<0.001) and the lowest mean intensities to 10-Hz burst. This rapid response to 100 Hz ICMS aligns with prior findings^13–15^ indicating robust initial firing followed by adaptation. Notably, there were no significant differences in preferential responses between excitatory and inhibitory subpopulations during the first 3 s (*p*>0.05).

Given that neural activity stabilizes after 10-20 s of ICMS, we next evaluated the last 10 s of ICMS to understand how individual neurons responded during the stable period. During the last 10 s, there was considerable variability in preferential responses. Some neurons had clearly higher activation by one train over the others. For example, we present an excitatory neuron with stronger peak activity in the last 10 s of ICMS for 10-Hz Burst (Fig. 3A, top) and an inhibitory neuron with stronger activity for TBS (Fig. 3A, bottom). However, most neurons exhibited only minor differences across the four presented trains. Among excitatory neurons, 34% preferred 10-Hz Burst stimulation, and 35% preferred TBS, although differences across trains were not statistically significant (*p*=0.14, Fig. 3D). The mean response in the last 10 s across all excitatory neurons was also significantly lower for 100-Hz train (0.61±0.04) than all other trains (*p*<0.05), while 10-Hz Uniform (0.83±0.05), 10-Hz Burst (0.79±0.05), and TBS (0.81±0.04) trains evoked similar mean intensities. Inhibitory neurons exhibited the highest response to TBS (*p*<0.001, Fig. 3E), and the percentage preferring TBS was significantly higher than in excitatory neurons (*p*<0.01, Fig. 3F). Interestingly, only 14% of inhibitory neurons responded most strongly to 10-Hz Burst, notably less than the 34% of excitatory neurons preferring this stimulus, though this difference was not quite significant (*p*=0.088).

Overall, 100-Hz ICMS evoked the strongest activation in the first few seconds of ICMS for all subtypes, followed by adaptation, leading to varied responses in the last 10 s. This strong but transient response may be expected as 100-Hz trains inject 10 times the stimulation pulses as the other trains tested in this study. Low-frequency trains sustained activation over long durations of ICMS, resulting in stronger activation during the last 10 s compared to the 100-Hz train (Fig. 3A-C). Interestingly, 63% of inhibitory neurons preferred TBS, while 35% of excitatory neurons did; conversely, 34% of excitatory neurons preferred 10-Hz Burst trains compared to 14% of inhibitory neurons. These results underscore that temporal patterns can increase or decrease sustained activation of individual neurons and may preferentially recruit different neuronal subpopulations, which could be pivotal for clinical therapies requiring activation of specific neural subpopulations.

### Individual excitatory neurons were more likely to adapt to ICMS, but individual responses were diverse

We showed that inhibitory neurons generally had facilitated ICMS responses, particularly for TBS. Nevertheless, all subpopulations exhibited variability in preference for different 30-s ICMS trains (Fig. 3D-E). We aimed to determine if diverse responses could be categorized based on temporal response patterns, expanding on previous classifications^13–15^ (Fig. 4). Neurons were classified based on changes in intensity over time: ‘facilitated’ (increased across the train), ‘adapted’ (initially increased, then decreased), ‘non-adapted’ (increased and remained stable), or decreased (dropped below baseline activity) to ICMS (Fig. 4A). ‘Adapted’ neurons were further divided based on the speed of adaptation: ‘rapidly adapting’ (RA) or ‘slowly adapting’ (SA).

**Figure 4:**
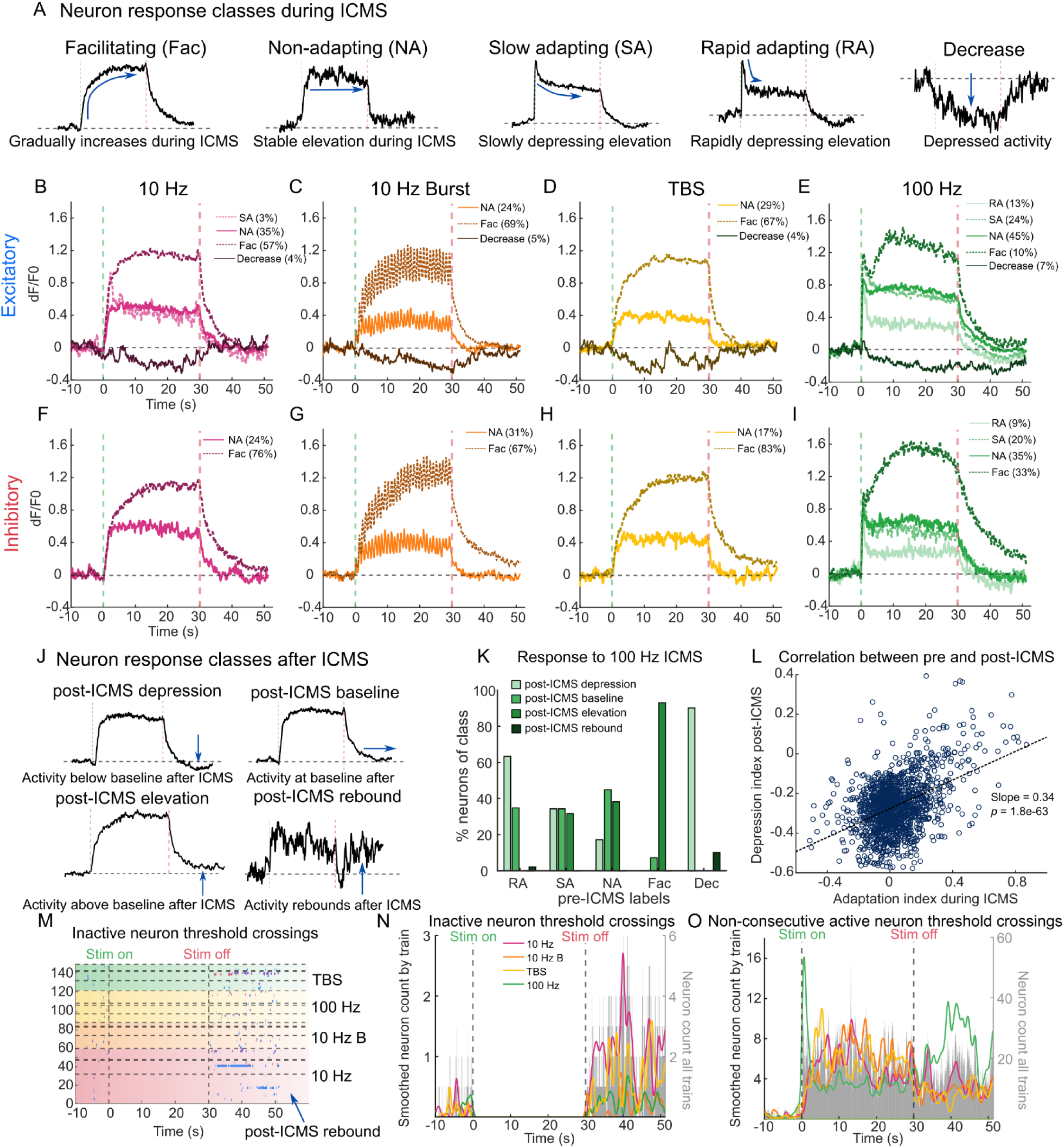
Individual inhibitory neurons adapted less and facilitated more to ICMS than individual excitatory neurons, which also correlated with less post-ICMS depression. A) Example traces showing ICMS-evoked response classes. B-M) Average responses of each class for excitatory (B-E), and inhibitory (F-I) neurons with each of the four ICMS trains. Inhibitory neurons generally exhibit more facilitating neurons, although all subpopulations contained diverse response types. J) Example traces showing post-ICMS response classes. K) ICMS-adapted neurons (RA, SA, Dec) were more likely post-ICMS depressed, ICMS-stable neurons (NA) were more likely post-ICMS stable, and ICMS-facilitated neurons (Fac) were more likely post-ICMS elevated. L) Strong correlation between ICMS-induced adaptation index with the post-ICMS adaptation index. The linear regression fit (black) and horizontal line marking zero adaptation (grey) are shown. M) Raster plot showing threshold crossing for inactive neurons, with some post-ICMS rebound. N-O) Post-stimulus time histograms for inactive neurons (N) and non-consecutive active neurons (O). Bars represent summed threshold crossings of all neurons across trains. Colored plots represent threshold crossings for individual trains smoothed with a 1 s Gaussian filter.

Low-frequency trains (10 Hz, 10-Hz Burst, TBS) predominantly evoked ‘non-adapting’ or ‘facilitating’ activity (Fig. 4B-D, F-H). As predicted from the average responses (Fig. 2C-F), a larger percentage of inhibitory neurons were ‘facilitating’ (67-83%) to low-frequency trains compared to excitatory neurons (57-69%) (Fig. 4B-H). Interestingly, neurons with decreased responses were only observed for excitatory and uncertain populations for every train (Fig. 4B-E, SFig. 3A-D). The TBS train facilitated the most inhibitory neurons (83%), while 10-Hz Burst trains facilitated the most excitatory neurons (69%), although these differences were not significant (*p*>0.05). Inhibitory neurons generally showed stronger facilitation to all trains than excitatory neurons, as quantified with the adaptation index (Fig. 2I). A smaller percent of neurons displayed facilitating responses to 100 Hz compared to low frequency trains (Fig. 4E,I). Like the other trains, inhibitory neurons had more facilitating neurons (33%) than excitatory (10%) to the 100-Hz train (*p*<0.05). Together, individual neurons exhibited different rates of adaptation based on the temporal pattern, with a higher percentage of inhibitory neurons facilitating compared to excitatory for all trains except the 10-Hz Burst.

Individual neurons show varied responses to the same ICMS parameters, but it is unclear if this variability reflects a stereotyped response of the individual neuron or is directly influenced by the applied temporal pattern. Neurons generally maintained the same classification across trains for the low-frequency trains (SFig. 4). Inhibitory neurons exhibited greater consistency across trains compared to excitatory neurons, with 66% classified as facilitating across all low-frequency trains, versus 55% for excitatory neurons. The 100-Hz train evoked the most diverse responses. While temporal patterns strongly influence individual neuron responses, consistent individual neuron responses across trains suggests a role for intrinsic neuron properties.

Overall, all subpopulations exhibited diverse responses to ICMS. Inhibitory neurons showed more consistent responses across trains and had a higher percent of facilitating neurons relative to excitatory neurons across all ICMS trains except for the 10-Hz Burst. The variability in individual neurons within each subpopulation may stem from factors such as proximity to the electrode, synaptic inputs, or variations in individual neuron morphologies^13,17^. These insights underscore the importance of understanding how ICMS affects both individual neurons and subpopulations, which is crucial for designing stimulation therapies aimed at continuously recruiting specific neuronal groups over time.

### Adaptation during ICMS correlated with depression post-ICMS

Previous work found that neurons adapt during ICMS and depress after ICMS^12,14,15^. However, it was unclear how adaptation during ICMS contributes to post-ICMS depression. Post-ICMS depression was evaluated for excitatory and inhibitory neurons (Fig. 4J, SFig. 3E-G), and its relationship to adaptation during ICMS was analyzed (Fig. 4K-L). Neurons were categorized based on their activity: below baseline (post-ICMS depression), decreased back to baseline (post-ICMS baseline), elevated to baseline (post-ICMS elevation), or if there was a distinct offset/rebound excitation (post-ICMS rebound) (Fig. 4J). Most neurons had activity equal to or greater than baseline following ICMS, especially with low-frequency trains (SFig. 3G). A smaller percentage of inhibitory neurons experienced post-ICMS depression (0-22%) than excitatory neurons (8-30%) (SFig. 3G). Additionally, higher frequencies evoked more post-ICMS depression (22-30%) than the lower frequency trains (0-16%) in all subpopulations. A small portion of neurons (0-11%) had post-ICMS rebound, typically having low activity during ICMS. TBS induced no post-ICMS depression in inhibitory neurons and the least post-ICMS depression in excitatory and uncertain subpopulations (6-8%) (SFig. 3G). In addition to less depression, inhibitory neurons had more elevated activity (48-69%) post-ICMS than excitatory neurons (36-46%) (SFig. 3G). Overall, excitatory neurons consistently exhibited more post-ICMS depression. TBS trains evoked the least post-ICMS depression across all subpopulations.

Adaptation during ICMS may indicate synaptic plasticity or neuronal exhaustion, leading to decreased activity post-ICMS^14,15,46^. Previous work did not clarify if post-ICMS depression is directly related to adaptation during ICMS. We evaluated this relationship and found that a larger percent of neurons classified as adapting during ICMS (RA or SA) were also classified as post-ICMS depressed (Fig. 4K). Specifically, neurons classified as RA during ICMS were more likely to be depressed post-ICMS (64%) than returned to baseline (36%) or elevated post-ICMS (0%). Conversely, the majority of ‘facilitated’ neurons had post-ICMS elevation (91%) and few had post-ICMS depression (<2%). Interestingly, neurons with decreased responses had the highest representation of post-ICMS rebound, indicating that rebound likely occurred due to release from inhibition (Fig. 4K). These relationships were further quantified by plotting the adaptation index during ICMS against the depression index after ICMS, revealing a highly significant correlation (*p*<0.001, Fig. 4L). These findings highlight the lasting effects of stimulation on the system and how the response to ICMS dictates the response and recovery of activity following ICMS.

### Adapting activity during ICMS may lead to disinhibition post-ICMS and stronger rebound activity

Neurons with decreased responses during ICMS were more likely to exhibit post-ICMS rebound. This prompted an investigation into whether inactive or non-consecutively active neurons would show similar rebound responses. Raster plots and post-stimulus histograms for inactive neurons revealed a clear increase in activity following ICMS (Fig. 4M-N). 10-Hz ICMS trials contained the largest number of inactive neurons and drove the strongest post-ICMS response. Non-consecutively active neurons exhibited rapid and strong adaptation following a brief activation during the initial seconds of 100-Hz ICMS, correlating with robust post-ICMS rebound (Fig. 4O). Overall, neurons with strongly adapted or decreased responses during ICMS were more likely to exhibit rebound responses, potentially indicating disinhibition.

### Inhibitory neurons are relatively more active farther from the electrode

We demonstrated that individual neurons exhibited varied degrees of adaptation during and depression after ICMS, with inhibitory neurons demonstrating overall lower likelihood of adaptation/depression. Previous studies indicate that neuronal responses to ICMS are influenced by proximity to the electrode^12–15^, potentially suggesting increased inhibitory influence in indirectly recruited populations^13,15^. To examine this, we analyzed if the activity and adaptation of each subpopulation varied with distance from the electrode, focusing on 100-Hz trains which exhibited more varied responses across neurons, similar to other trains (SFig. 5). There was no difference in the average distance of active excitatory and inhibitory neurons from the electrode (*p*=0.81, Fig. 5A). Active neuron counts were evaluated in 25 µm bins from the electrode (Fig. 5B). For excitatory neuron, active counts peaked around 100-125 µm from the electrode, while active inhibitory neuron counts peaked around 175-200 µm. Beyond 325 µm, no continuous active inhibitory neurons were detected, although non-consecutive active inhibitory neurons were observed. Inhibitory neurons exhibited a narrower range of continuous activation (50-325 µm) compared to excitatory neurons (0-525 µm), possibly influenced by the smaller sample size of inhibitory neurons.

**Figure 5:**
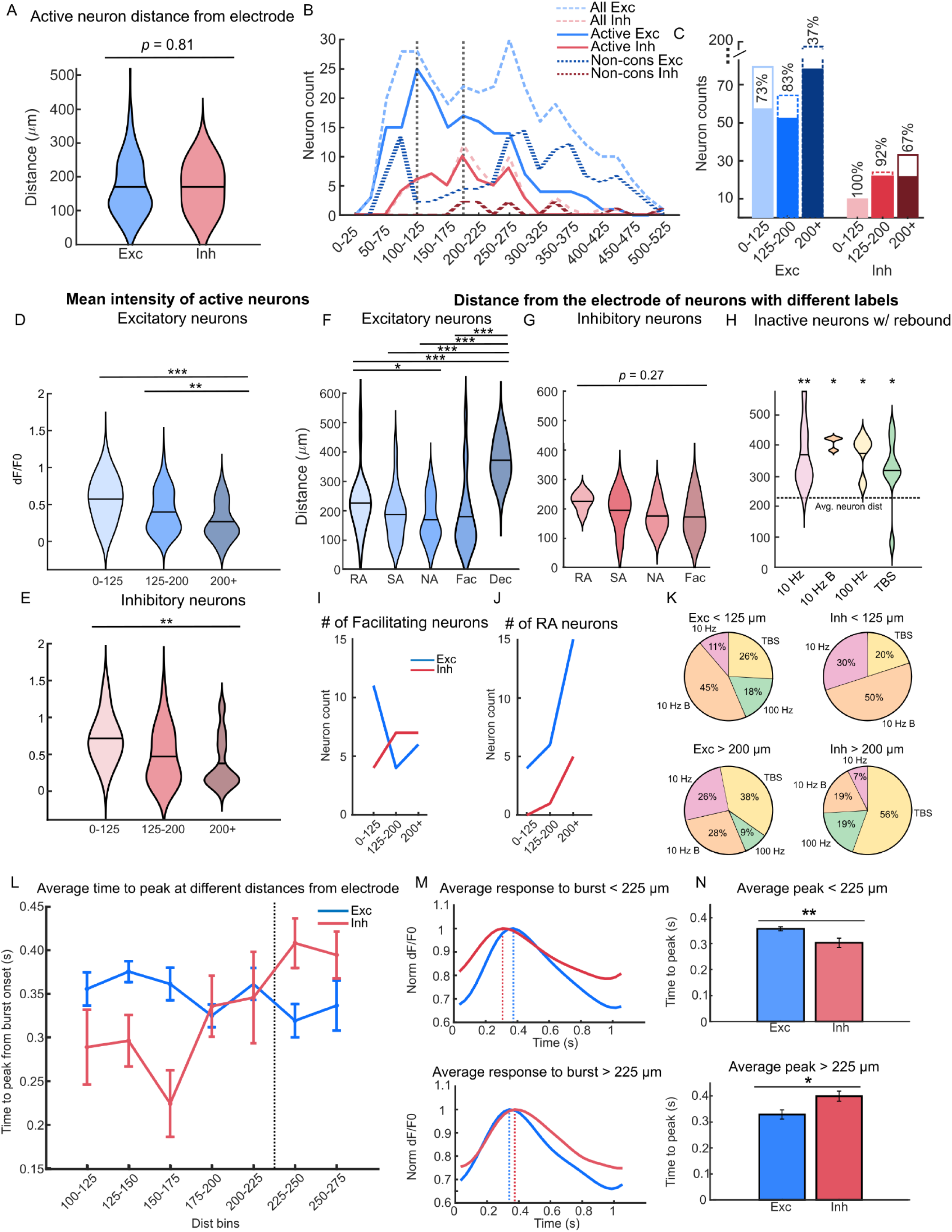
Inhibitory neurons are more active farther from the electrode, and farther facilitating neurons are more likely to be inhibitory. A) Violin plots show no significant difference in average distance from the electrode between active excitatory and inhibitory neurons. B) Distribution of excitatory and inhibitory neurons across 25 µm bins away from the electrode. Active excitatory neurons peak around 100-125 µm while active inhibitory neurons peak around 175-200 µm. C) The number of excitatory and inhibitory neurons in 0-125, 125-200, and 200+ distance bins. The dotted bar shows all neurons while the solid filled bar shows only active neurons. The y-axis is discontinuous between 80-200 for illustration. D-E) Violin plots of the mean intensity of excitatory (D) and inhibitory (E) neurons. Intensity decreases for both excitatory and inhibitory neurons farther from the electrode. F-G) Violin plots of the average distance for ICMS-evoked classes for excitatory (F) and inhibitory neurons (G). RA neurons are farther and NA/Facilitating neurons are closer from the electrode. H) Violin plot for distance of inactive excitatory neurons with rebound. The horizontal line marks the average active neuron distance from the electrode (234 µm). All violin plots are above the average, indicating inactive-rebound neurons are farther from the electrode. I) Facilitating neurons farther from the electrode are more likely to be inhibitory. J) Neurons farther from the electrode are more likely to be RA for both excitatory and inhibitory. K) Pie plots showing neurons closer to the electrode (top row) prefer 10-Hz Burst and neurons farther from the electrode (bottom row) prefer TBS. L) Line plot showing average time-to-peak after burst onset for excitatory and inhibitory neurons in 25 µm bins. Inhibitory neurons have faster times-to-peak <225 µm from the electrode, but slower times to peak >225 µm. The vertical line marks the flip at 225 µm. Error bars indicate standard error across neurons within each bin. M) Average traces of last six bursts for excitatory and inhibitory neurons that are <225 µm (top) or >225 µm (bottom). N) Bar plots showing the average time-to-peak after burst onset for exc vs inh neurons that are <225 µm (top) or >225 µm (bottom). Error bars indicate standard error across neurons within each bin.

To compare how neuron activation changed across distance, neurons were split into distance bins (0-125, 125- 200, and 200+ µm) with roughly equal numbers of active excitatory neurons, ensuring sufficient statistical power (Fig. 5B). The number of active inhibitory neurons doubled from the 0-125 µm bin to the 125-200 µm bin and was stable between the 125-200 and 200+ µm bins (Fig. 5C). In the 200+ µm bin, a lower percentage of excitatory neurons were active overall (37%) compared to inhibitory neurons (67%), suggesting that indirect recruitment more strongly activates inhibitory neurons, aligning with previous optogenetic results^47,48^. Next, the mean intensity of neurons was evaluated in each bin. Both excitatory and inhibitory neurons decreased in activation intensity farther from the electrode (*p*<0.01, Fig. 5D,E). Together, these results indicate that excitatory neurons are more active near the electrode, while inhibitory neurons are more active farther from the electrode, agreeing with previous results using shorter trains^17^. Overall, these findings highlight that individual neuron responses correlate with their distance from the electrode, with increasing inhibitory drive farther from the electrode. This insight may elucidate phenomena such as adaptation of neurons distant from the electrode and robust excitatory recruitment near it.

### Excitatory neurons showed adaptation and post-ICMS rebound farther from the electrode, while inhibitory neurons were more stable across space

We showed that neurons within each subpopulation exhibited varied responses (adapting or facilitating) to ICMS. Previous studies showed these responses correlate with distance from the electrode^13–15^, but the influence of inhibitory neurons on these distance relationships remain unclear. Consistent with previous findings^13^, neurons classified as RA were typically located farther from the electrode compared to NA neurons (Fig. 5F,G, SFig. 5E-G). Additionally, neurons with decreased responses were significantly farther from the electrode than other groups, suggesting synaptic effects (Fig. 5F, SFig. 5E,G). Given that these decreased responses were observed farther from the electrode, we investigated whether inactive neurons with post-ICMS rebound responses were farther from the electrode (Fig. 5H). Indeed, inactive neurons showing rebound responses were on average farther (300-400 µm) from the electrode than the average neuron (Fig. 5H, *p*<0.05). RA neurons being situated farther from the electrode could suggest stronger synaptic inhibition, which may be driven by increasing inhibitory drive. Indeed, inhibitory neurons farther from the electrode are more likely to facilitate to ICMS than excitatory neurons (Fig. 5I). Facilitating neurons >125 µm from the electrode — and therefore more likely indirectly recruited — were predominantly inhibitory, despite ∼3x more active excitatory neurons in this range (Fig. 5I). This was not true for RA neurons, which were mostly excitatory (Fig. 5J). Together, these results suggest that putative direct activation results in more stable, stereotyped responses, while putative indirect activation results in more complex effects, such as facilitation of inhibitory neurons, decreased activation of excitatory neurons, and rebound responses due to disinhibition.

### Neurons closer to the electrode preferred 10 Hz Burst trains, while neurons farther preferred TBS

Given differences in the relative activation of excitatory and inhibitory neurons based on distance and the preference of TBS and 10-Hz Burst trains for these subpopulations, we examined whether neurons had preferences for TBS or 10-Hz Burst based on their electrode proximity. Neurons <125 µm from the electrode more likely preferred 10-Hz Burst in both subpopulations (45-50%, Fig 5K). Conversely, both subpopulations were more likely to prefer TBS farther from the electrode (38-56%). This suggests that TBS may effectively drive activation through synaptic mechanisms, enhancing inhibitory drive in the network over time (Fig. 5C,I). In contrast, 10-Hz Burst appears more effective at stimulating peak activity in directly recruited neurons, which predominantly include excitatory neurons. Overall, TBS demonstrated superior efficacy in activating neurons farther from the electrode, suggesting TBS enhances inhibitory drive in the network by facilitating synaptic recruitment of inhibitory neurons.

### Inhibitory peaked faster than excitatory neurons near the electrode, but peaked slower farther away

In feedforward inhibition, inhibitory neurons are recruited faster or simultaneously with excitatory neurons, whereas in feedback inhibition, inhibitory neurons are recruited afterwards^24,49,50^. We aimed to determine if similar activation delays exist between excitatory and inhibitory neurons during ICMS. Analysis of 10-Hz Burst trains revealed distinct neuronal modulation with each burst, enabling comparison of time-to-peak activation between subpopulations (Fig. 5L-N). Inhibitory neurons generally showed faster times-to-peak than excitatory neurons closer to the electrode, with a reversal of this trend beyond 225 µm (Fig. 5L). When divided into bins (<225 µm and ≥225 µm), inhibitory neurons within <225 µm exhibited faster time-to-peaks than excitatory neurons (*p*<0.01, Fig. 5L,M top). Beyond 225 µm, inhibitory burst responses tended to have slower times-to-peaks than excitatory responses, though this difference was not statistically significant (*p*=0.07, Fig. 5L,M, bottom). Morphological differences ̶ such as shorter processes, myelination, and faster channel dynamics ̶ may contribute to disparities in putative direct recruitment. Circuit-level mechanisms, such as feedback inhibition, could influence differences in putative indirect recruitment. This finding highlights important spatiotemporal differences in the recruitment of excitatory and inhibitory neurons which informs our understanding of cortical circuits as well as ICMS propagation in the network.

### Neuron-neuron and neuron-neuropil correlations are strongest for ICMS near the electrode, but strongest for baseline activity farther

We showed that inhibitory neurons exhibited higher activity and are more likely to facilitate farther from the electrode than excitatory neurons. Additionally, both excitatory and inhibitory neurons showed higher rates of adaptation (RA or SA) farther from the electrode. We hypothesized that synaptically facilitated inhibitory neurons activated by ICMS may inhibit neighboring neurons, leading to adaptation farther from the electrode. However, proximity between neurons may indicate they are part of the same functional unit, raising questions about correlations between inhibitory and nearby excitatory neurons. Correlations in ICMS-evoked and baseline activity were assessed for individual active inhibitory neurons and all other active neurons, as well as individual excitatory neurons and all other active neurons (Fig. 6A). Generally, neurons closer to the electrode (<200 µm) exhibited higher correlations in ICMS-evoked activity than those farther away (>200 µm, Fig. 6B-D). Previous studies using iGluSNFR suggest that the direct activation volume is <100 µm from the electrode at 20 µA^15^, leading to antidromic activation of neuronal soma in this range. Therefore, neurons within 100 µm are likely to have high correlations due to their direct entrainment to the stimulus profile. Conversely, neurons indirectly recruited may show greater variations in responses due to synaptic integration, resulting in lower correlation values that decrease over distance. All subpopulations had higher correlations between neurons near each other (*p*<0.01, Fig. 6B-D). Overall, neurons closer together, regardless of subpopulation, are more likely to exhibit correlated responses to ICMS, with stronger correlations observed near the electrode compared to farther distances.

**Figure 6:**
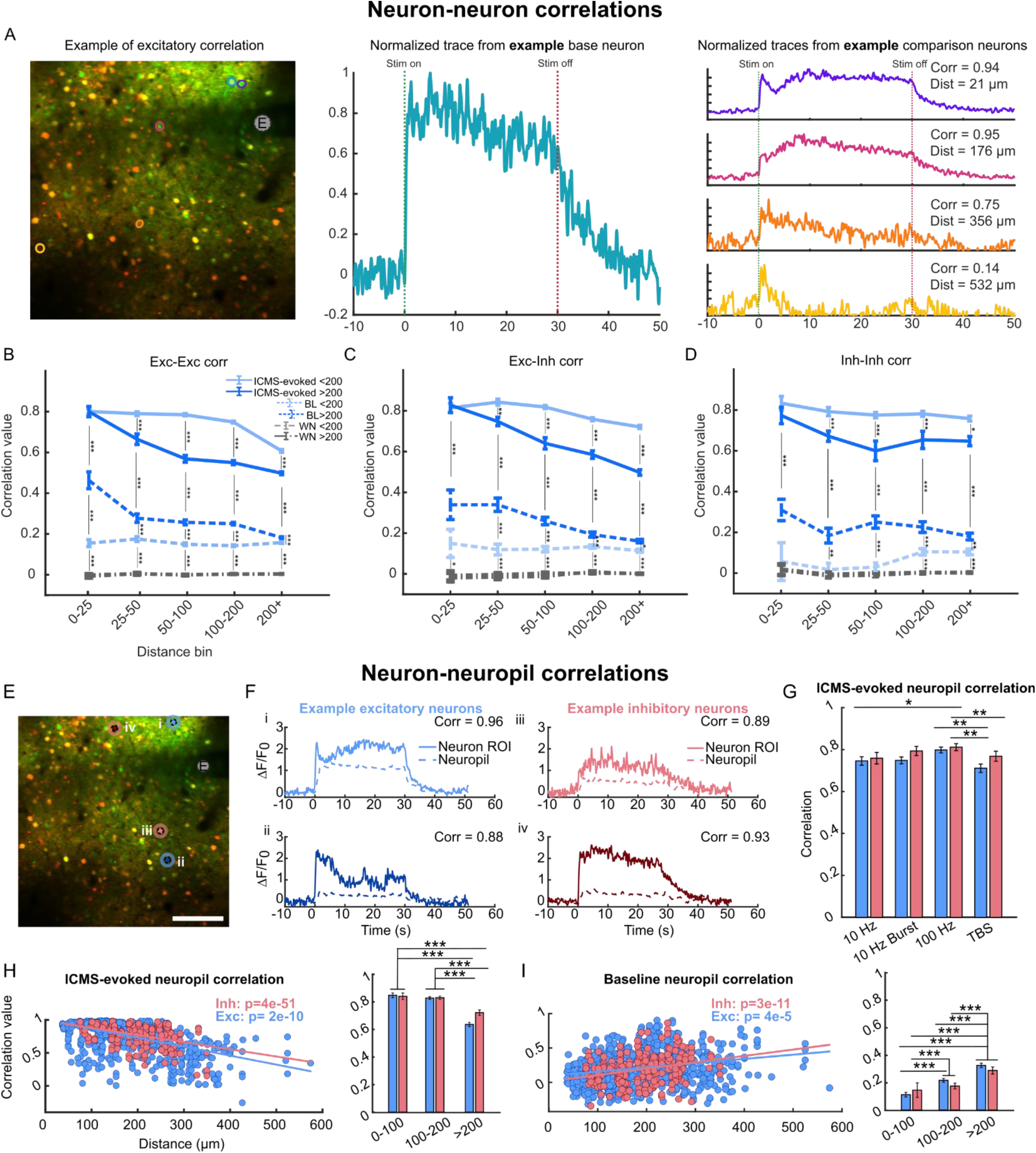
Neurons closer to the electrode had stronger neuron-neuron and neuron-neuropil correlations during ICMS but weaker correlated activity during baseline compared to neurons farther from the electrode. A) Example image showing four individual neurons of different distances correlated in activity to a single neuron. Note that neurons farther from the single neuron have lower correlations. B-E) Average correlation value of neurons within different distance bins from the electrode for B) excitatory-excitatory, C) excitatory-inhibitory, and D) inhibitory-inhibitory correlations. Neurons closer together tend to have higher correlations. ICMS-evoked responses have higher correlations than baseline responses. Error bars indicate the standard error across all calculated correlations. BL represents correlation to baseline activity (15 s pre-ICMS) while WN represents correlation to white noise processed with the same filter. E) Example image with four neurons and the area of neuropil surrounding each neuron. F) Traces for selected neuron responses to ICMS and the corresponding activation of the surrounding neuropil. G) Bar plot showing the neuron-neuropil correlation for excitatory and inhibitory subpopulations for each ICMS train. Error bars indicate standard error across neurons. H) Scatter plots with linear fits showing individual ICMS-evoked neuron-neuropil correlations as a function of distance showing correlations decreased with increasing distance. I) Scatter plots with linear fits showing individual baseline neuron-neuropil correlations as a function of distance show increased farther from the electrode.

During ICMS, neurons exhibited higher correlation values compared to baseline across all subtypes and comparisons (*p*<0.001, Fig. 6B-D), which is consistent with ICMS inducing stronger and focal activation relative to sensory evoked responses^51–53^. Baseline correlations exhibited significantly higher correlations than white noise (*p*<0.05, Fig. 6B-D) except for inhibitory-inhibitory correlations <100 µm from the electrode (Fig. 6D). Notably, neurons near the electrode displayed *lower* baseline correlations with each other, while those farther from the electrode showed *higher* baseline correlations. Lower correlation for neurons near the electrode suggests that electrode insertion and/or strong ICMS activation may have disrupted the local network. It is likely that electrode insertion itself caused damage to the surrounding circuitry and connections among neurons, resulting in low baseline activity correlations. Despite this disruption, neurons near the electrode exhibited high correlations during ICMS due to direct recruitment and entrainment to the stimulation profile. These findings underscore significant differences between normal cortical activity and ICMS-evoked responses, which are influenced by both tissue damage from electrode insertion/stimulation and the nature of direct versus indirect neural activation profiles.

While neurons can be individually investigated using two-photon imaging, background neuropil activity may also contain important information. Neuropil activity is influenced by neuronal processes (such as dendrites and axons) and activity from neurons outside the imaging plane, resembling local field potential (LFP) activity^54^. For ICMS evoked activity, neuropil is thought to largely reflect the activity of directly recruited axons, offering insights into how closely a neuron follows the activation pattern of directly activated neurons^11,14,15^. To elucidate influences on excitatory versus inhibitory neuron activity across distances and their correlations with neuropil activity, we examined whether excitatory or inhibitory neurons exhibited stronger correlations with neuropil within an annular region surrounding activated neurons (Fig. 6E,F).

Excitatory and inhibitory neurons showed similar correlations with the neuropil across different temporal patterns (Fig. 6G). However, inhibitory neurons exhibited slightly higher correlations on all trains but especially for TBS, although this difference was not significant (*p*=0.064). Distinct ICMS trains influenced neuron-neuropil correlation differently, with 100 Hz producing the highest correlation and TBS producing the lowest (*p*<0.01, Fig. 6G). Similar to neuron-neuron correlations, neuron-neuropil correlation decreased with distance from the electrode (*p*<0.001, Fig. 6H, SFig. 6). Neurons within 0-200 µm of the electrode showed high (>0.9) neuron-neuropil correlations (Fig. 6H, SFig. 6), suggesting direct activation. Baseline correlations were lower than ICMS-evoked correlations, and neuron-neuropil correlations increased with distance from the electrode (opposite to ICMS-evoked activity), suggesting electrode-induced disruption of the natural sensory circuit (Fig. 6I). Neuropil-neuron correlation mirrored neuron-neuron correlations, highlighting similar relationships without further revealing distinct differences in excitatory versus inhibitory recruitment relative to directly activated processes.

Overall, neurons closer together are generally more correlated to both ICMS evoked responses and baseline sensory activity. Due to the synchronous activity evoked by ICMS, all neuron subpopulations showed highly correlated neuron-neuron and neuron-neuropil activity during ICMS. Neurons closer to the electrode, likely recruited directly, maintain high ICMS-evoked correlations that remain stable over distance until beyond the direct activation range. Conversely, neurons farther from the electrode exhibit sharper declines in neuron-neuron correlation with increasing distance. Proximity suggests shared synaptic inputs among closely spaced neurons, enhancing correlated activity. Previous studies noted stronger firing correlations among closely located neurons for both excitatory and inhibitory subtypes^55,56^. Neurons closer to the electrode, however, display lower baseline neuron-neuron and neuron-neuropil correlations, likely due to electrode interference with sensory circuits, yielding consistent low correlations irrespective of distance. Nonetheless, ICMS effectively synchronizes these neurons. These findings underscore crucial distinctions in direct versus indirect ICMS recruitment, impact of electrode insertion on sensory circuits, and differential responses of excitatory and inhibitory neurons to ICMS and baseline cortical activity.

## Discussion

### Inhibitory neurons exhibit significantly more facilitation to prolonged ICMS than excitatory neurons

We identified significant differences in inhibitory and excitatory neuron responses to long bouts of ICMS. Crucially, many of these observed differences occurred after 10 s of continuous ICMS and were therefore dependent on long-duration ICMS, unlike previous work^17^. VGAT-labeled inhibitory neurons exhibited greater recruitment, mean intensity, and facilitation compared to non-VGAT labeled excitatory neurons (Fig. 2). Additionally, facilitating neurons farther from the electrode were much more likely to be inhibitory neurons (Fig. 6). This evidence suggests that synaptically recruited inhibitory neurons are more strongly activated over time by all ICMS trains, which may drive depression of the network, observed here and previously^12–15^. However, we could not directly confirm in our results that the increase in inhibitory neuron activity directly caused adaptation of excitatory neurons, as connections between neurons could not be imaged. Notably, during 100 Hz ICMS, 33% of inhibitory neurons were facilitating, contrasting with only 13% of excitatory neurons (Fig. 5). Still, 31% of inhibitory neurons showed adaptation to 100-Hz trains, suggesting inhibitory-inhibitory interactions. Inhibitory-inhibitory connections can also drive disinhibition of excitatory neurons, resulting in facilitation or rebound responses in excitatory neurons (Fig. 4). These complex interactions of excitatory and inhibitory neurons, and particularly the strong facilitation of inhibitory neurons, was not observed previously with shorter durations of ICMS^17^. Here, long durations of ICMS facilitated a significant portion of indirectly activated inhibitory neurons, which can drive complex responses in the network based on synaptic connections, such as inhibition and disinhibition.

### Burst-paradigms drive stronger activity over time, with inhibitory neurons preferring TBS and excitatory neurons preferring 10-Hz Burst

We tested four temporal patterns and found that the high-frequency train (100 Hz) evoked the strongest activity in the first few seconds, but ultimately resulted in stronger adaptation across 30 s, likely due to either an increase in inhibitory drive or metabolic exhaustion^32^ of the recruited populations (Fig. 7D). In contrast, low-frequency ICMS produces more sustained but weaker activation (Fig. 7E). For sensory restoration, low-frequency ICMS produces inconsistent percepts^2^; evoked sensations feel like naturalistic “taps” on some electrodes but are imperceptible on other electrodes^2,57^. Together, high and low-frequency ICMS are not ideal for evoking strong and consistent activity and perception over time.

**Fig. 7:**
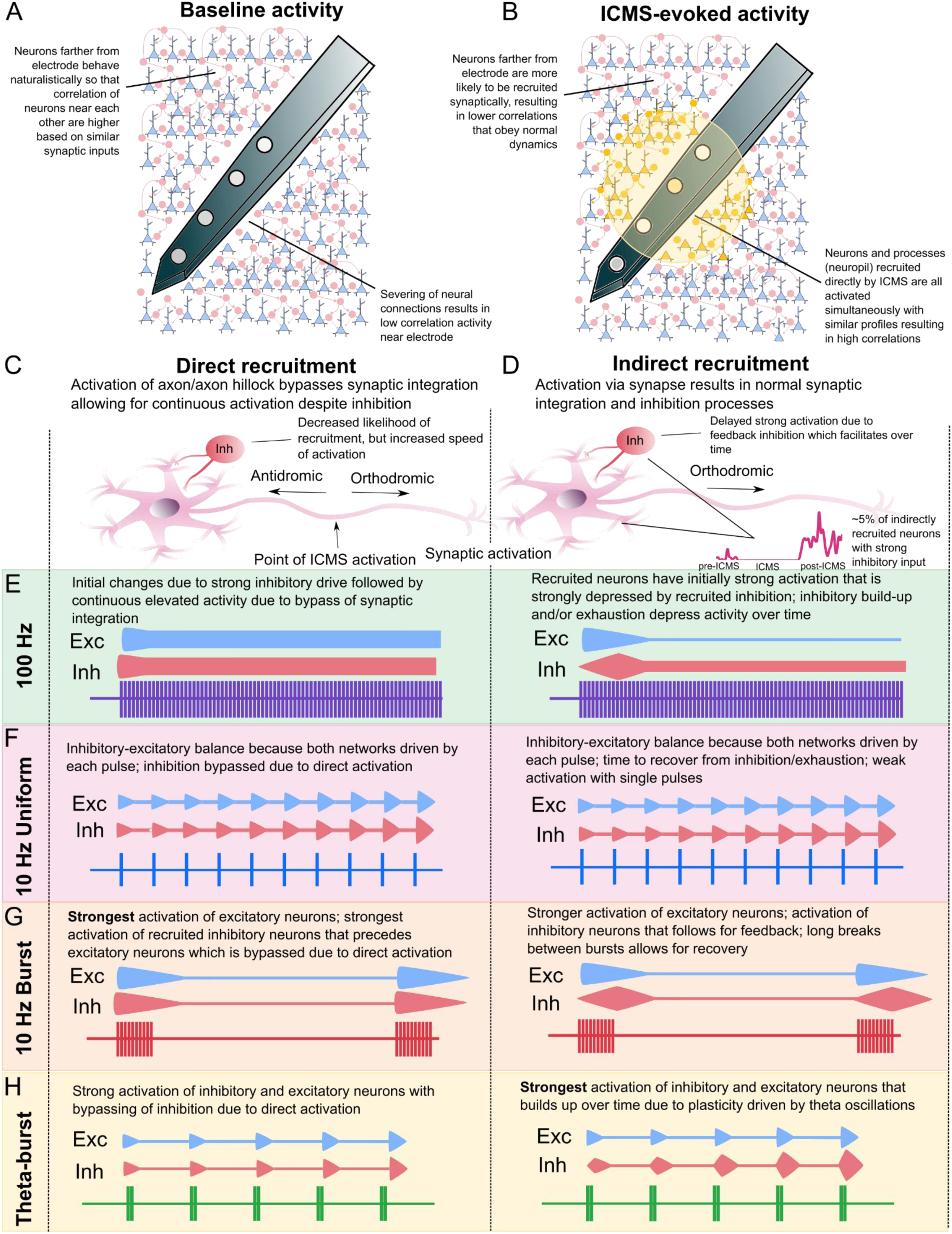
Explanatory framework. A) The electrode disrupts nearby neuron connections, reducing baseline correlation; farther neurons maintain higher correlations that drop-off with distance. B) Direct recruitment near the electrode, especially of excitatory neurons due to morphology, results in high correlations during ICMS. Indirect recruitment results in lower correlations that drop-off with distance due variance in synaptic integration. C) Neurons near the electrode are primarily recruited directly at the axon/axon hillock. Direct activation bypasses normal synaptic processes by recruiting the axon, decreasing inhibitory influence. D) Neurons farther from the electrode are primarily recruited indirectly via synaptic integration in the soma. Some neurons recruited indirectly are inhibited, which results in a decrease in activity during ICMS often followed by post-ICMS rebound. E) 100-Hz/high-frequency trains induce rapid adaptation in recruited neurons. Neurons near the electrode to have stable recruitment after initial adaptation due to bypassing of synaptic integration (left). Farther neurons undergo continual adaptation due to inhibitory drive and exhaustion. F) 10-Hz Uniform trains weakly recruit neurons with single pulses and the space between pulses allows for recovery from inhibition. G) 10-Hz Burst trains strongly activate excitatory and inhibitory neurons near the electrode, with inter-burst intervals allowing sufficient time for recovery. H) TBS drives strong activation of neurons farther from the electrode (right) over time, with rhythmic oscillations produced by TBS facilitating neurons (esp. inhibitory).

Burst-type paradigms mimic physiologic firing patterns, offering intense activation interspersed with recovery phases. Burst paradigms have been used successfully to maintain perception over longer periods of times in humans, although can extinguish percepts across 60 s of ICMS^28^. Here, the two bursting paradigms resulted in preferential activation of excitatory or inhibitory populations (Fig. 4). For 10-Hz Burst trains, 34% of excitatory neurons had the strongest peak activation, compared to only 14% of inhibitory neurons (Fig. 3). Additionally, all subpopulations near the electrode exhibited a preference for 10-Hz Burst (Fig. 5K). However, during the off-phase of the burst cycle, excitatory neuron activity decreased to levels lower than those seen with other low-frequency trains, leading to an overall reduction in average excitatory activity during the initial stimulation period (Fig. 3B). Together, 10-Hz burst stimulation initially activates excitatory neurons the least compared to other low-frequency trains, but by the end of the stimulation period, excitatory neurons respond more robustly.

TBS preferentially activates farther neurons, suggesting enhanced synaptic facilitation. TBS has been used previously in TMS and other stimulation paradigms^58–60^. TBS mimics theta rhythms in the brain, which are dependent on inhibitory neurons and associated with synaptic plasticity^61–64^. TBS may then be more successful in driving synaptic facilitation in the network by inducing oscillations following the frequency of stimulus delivery^65^. Here, TBS led to significantly stronger activation of inhibitory subpopulations, with 63% of inhibitory neurons have the strongest response in the last 10 s compared to 34% of excitatory neurons (Fig. 4). Additionally, neurons farther from the electrode were more likely to prefer TBS (Fig. 5K). These results suggest that rhythmic theta activation may synaptically facilitate neurons, with synaptic recruitment preferentially activating inhibitory neurons.

Overall, burst-type paradigms are better at maintaining strong activation over time, with different burst patterns producing preferential activation of different populations of neurons. Burst-type paradigms share similarities to “biomimetic” or “dynamic” paradigms used recently, which deliver bursts of high amplitude/frequency ICMS during important transient sensory events^13,66–68^. Selective recruitment of excitatory neurons may be desirable for certain applications (such as sensory restoration, motor rehabilitation, etc.) while selective recruitment of inhibitory neurons may be desirable in other applications (seizure control, schizophrenia, etc.). These insights underscore the importance of exploring tailored stimulation parameters to effectively modulate excitatory and inhibitory neuron dynamics for targeted therapeutic strategies.

### Putative directly recruited neurons have lower baseline correlations due to circuit disruption, but higher ICMS correlations due to strong simultaneous recruitment

Correlation in neural activity suggests synaptic connectivity^69,70^. Neurons closer to each other are more likely to be synaptically connected and receive similar synaptic input, resulting in higher baseline correlations as observed farther from the electrode (Fig. 6). We observed lower baseline correlations closer to the electrode, suggesting disruption of synaptic connections (Fig. 7A). Indeed, electrode insertion can lead to neural death and circuit changes near electrodes^71–74^. These damaged neurons are also more likely to be recruited directly by ICMS due to electrode proximity (Fig. 7B). Direct activation results in higher correlations of activity during ICMS, despite impaired synaptic connectivity. Inhibitory feedback may be less likely to impinge on neurons near electrodes because, 1) of disrupted synaptic connections, and 2) direct recruitment bypasses inhibitory integration at the soma (Fig. 7C). Additionally, inhibitory neurons may be less likely to be directly recruited by ICMS due to their morphology^17^. Because of these effects, all subpopulations near the electrode show more stable/fewer adapting responses to ICMS^12–15^. Farther from the electrode, by contrast, neurons have lower correlations that decrease with distance apart, as is normally observed in sensory cortices^55,56^ (Fig. 6).

### Indirect recruitment results in excitation, inhibition, or disinhibition based on synaptic inputs and applied temporal profile, with theta-burst stimulation driving the strongest synaptic excitation over time

Synaptic recruitment can drive more varied and complex responses, such as inhibition and disinhibition, based on converging glutamatergic and/or GABAergic input onto synapses (Fig. 7D). 100-Hz profiles induce continuous activity in directly recruited neurons, but strongly adapting activity in indirectly recruited neurons (Fig. 7E). Inhibition then likely plays a role in adaptation of activity observed over time, since directly-recruited neurons are less impacted by inhibitory feedback. However, it is also possible that for neurons near the electrode that ATP depletion due to strong recruitment quenches the ability for neurons to clear calcium, resulting in elevated Ca^+2^ intensity despite decreases in physiological firing^75^. The lower frequency and burst paradigms are able to maintain activity for both directly and indirectly recruited neurons due to interpulse/interburst spacing (Fig. 7F-H). Low frequency trains evoke weak but stable activation in the population (Fig. 7F). 10-Hz Burst drives strong peak activity in directly recruited excitatory neurons (Fig. 7G). TBS is able to drive strong activity through synaptic mechanisms, particularly in inhibitory neurons (Fig. 7H). This explanatory framework highlights the interplay of circuit disruptions driven by the electrode, mechanisms of neuron recruitment, and the effects of temporal parameters on activation patterns. These observations underscore the need for improved technologies for interfacing with the brain as well as further understanding and optimization of temporal profiles for targeted neural recruitment.

### Limitations

We sought to understand the impact of ICMS on inhibitory neurons and their effect on overall network activation. While our results have strong implications about direct and indirect activation, the temporal resolution of calcium responses (hundreds of milliseconds) is too slow to detect latencies between direct and indirect activation (<10 ms). Furthermore, calcium latencies limit the ability to measure entrainment of neurons to different temporal profiles as well as activity within the first few hundred milliseconds of ICMS or post-ICMS. Emerging technologies, like genetic encoded voltage indicators (GEVIs) with higher spatial resolution may reveal new insights into these dynamics^76,77^. Temporal patterning can produce varying effects in neurons, but the temporal profiles presented here were not completely exhaustive. Future work should explore other temporal patterns — as well as other stimulus parameters — for optimization of ICMS profiles in therapeutic applications. Separation of inhibitory and excitatory subpopulations was also not trivial here, making our classifications less certain. Using multiple wavelengths for measurement could allow for better separation. Mice were tested the day of the procedure after awaking from anesthesia. It is unclear then if residual anesthesia effects or acute surgical responses could impact these results. Future work should compare ICMS with anesthesia and under chronic conditions to better understand how these effects may impact the cortical response to ICMS.

## Conclusion

Overall, our findings reveal distinct microstimulation recruitment patterns between excitatory and inhibitory neurons, which is strongly dependent on distance from the electrode and the temporal pattern provided. Neurons near the electrode were more stable and stereotyped in activation, while neurons farther from the electrode had more varied and complex responses, such as inhibition and disinhibition. We found that inhibitory neurons are relatively more active farther from the electrode and are more likely to facilitate to ICMS over time. TBS was also more likely to drive strong activity in inhibitory neurons and neurons farther from the electrode. Based on correlation analysis, neurons near the electrode appeared to have disrupted synaptic inputs. Despite this, these neurons were the most strongly activated by ICMS, especially for the 10-Hz Burst profile. Together these results indicate that burst paradigms (similar in design to biomimetic/dynamic paradigms) are generally better at driving activity over long durations, but 10-Hz Burst might be preferred for directly activated excitatory neurons, while TBS may be better for indirectly activated inhibitory neurons. This work then reveals crucial insights into the mechanisms underlying network activation by ICMS with clinically relevant paradigms, and also suggests specific types of temporal patterns (particularly burst profiles) as a promising path towards improved clinical stimulation therapies.

## Acknowledgements

This work was supported by the National Institute of Neurological Disorders and Stroke of the National Institutes of Health under Award Numbers R01NS105691 and R01NS115707 and the National Science Foundation under award CAREER CBET 1943906. Christopher Hughes was supported by the NINDS T32 NS08674932 training award for the study of the neurobiology of neurological disease and the NIH BRAIN F32MH130022. Kevin Stieger was supported by the NIH NINDS F31NS125982.

## Methods

### Surgery, virus injection, and electrode implantation

Transgenic VGAT-Cre (n=4) mice (Jackson Laboratories, Bar Harbor, ME) were used in these experiments. All surgical interventions were performed on adult mice (>4 weeks of age) that are housed in social housing with 12-h light/12-h dark cycles and free access to food and water until experimentation. Weeks prior to electrode implantation, mice received intracortical viral injections of AAV-Syn-GCaMP7b for imaging of excitatory neurons and AAV-FLEX-tdTomato to identify VGAT-inhibitory neurons. Previous work has shown strong overlap with inhibitory neurons (>95%, Gad67, PV, SST) and little overlap with excitatory neurons (<1%, VGlut1) in cerebral cortex for VGAT-tdTomato^78^. Injections were completed with a micromanipulator, 5-10 um diameter glass pipette, and micro-injector (Toohey Company, Fairfield, NJ, USA). Anesthesia was induced with a mixture consisting of 75 mg/kg ketamine and 7 mg/kg xylazine administered intraperitoneally (IP) and updated with 40 mg/kg as needed, as described previously^14,15^. Head-fixed mice were given circular craniotomies (∼3 mm diameter) over visual cortex with a dental drill. Virus was bolus injected into the visual cortex under aseptic conditions. Following injection, the craniotomy was sealed with a glass cover slip and secured with dental cement. At least 3 weeks after viral injection, the mouse was anesthetized with isoflurane (concentration) and placed in a head-fixed apparatus. Once anesthetized, the cover glass was removed from the skull and the electrode was implanted. We used a single shank Michigan style array with 3-mm long shanks with 4 703 µm^2^ electrode sites spaced 100 µm apart (NeuroNexus Technologies, Ann Arbor, MI) in each animal. Electrodes were implanted at a 30° angle using an oil-hydraulic Microdrive (MO-81, Narishige, Japan). Electrode arrays were inserted to depths of 600 µm at a speed of 200 µm/s. Following insertion, the animal was taken off of isoflurane and left to wake up (based on behavioral indicators i.e. whisking, hindlimb movement) at which point the data collection began.

### Electrical stimulation

We used a TDT IZ2 stimulator controlled by an RZ5D system (Tucker-Davis Technologies, Alachua, FL) to apply current-controlled cathodic leading biphasic asymmetric (200 μS cathodic phase, 100 μS interphase, 400 μS anodic phase). This pulse shape was selected to match the previous human experiments^1,2^, as it has been shown that pulse shape itself can affect recruitment^11^. Pulse amplitude adjusted the cathodic and anodic phase of each pulse while pulse frequency adjusted the number of pulses presented in a single second of stimulation. We used amplitudes of 10 µA (2 nC per phase) and frequencies of 10 or 100 Hz. These limits were chosen based on established safety limits to reduce electrode and tissue damage that have been described previously^79–83^. Concurrent imaging and stimulation trials were acquired for 60-s (10 s pre-ICMS, 30 s ICMS, 20 s post-ICMS) with each parameter set. We stimulated one electrode within the imaging plane at amplitudes of 10 µA for 30 s with four different ICMS trains: 10 Hz Uniform, 100 Hz Uniform, 10-Hz Burst (10 pulses provided within 100 ms followed by 900 ms of no ICMS), and Theta Burst Stimulation (two pulses provided 10 ms apart every 200 ms). Trials were randomly interleaved to remove ordering effects.

### Two-photon imaging

A two-photon laser scanning microscope (Bruker, Madison, WI) and an OPO laser (Insight DS+; Spectra-Physics, Menlo Park, CA) tuned to a wavelength of 920 nm at 15 mW was used for the entirety of this study. A 16X 0.8 numerical aperture water immersion objective lens (Nikon Instruments, Melville, NY) was used for its 3- mm working distance. The imaging was in plane with the electrode site, 150–300 µm below the surface of the brain. To image neural dynamics during electrical stimulation, time series images were recorded at 30 Hz and 1× optical zoom. The imaging plane for the time series was set over the electrode contact so that the stimulating contact is visible during imaging session, as described previously^12,14,15^.

### Image processing

Cells were manually outlined using a 2D projection image of the standard deviation of fluorescence using ImageJ (NIH). For each outlined cell, the mean fluorescence activity of the cell was extracted over time. Analysis of the fluorescence data was conducted in MATLAB (Mathworks, Natick, MA). All fluorescence was filtered using a 15- point Gaussian temporal filter. Fluorescence was transformed into a measure normalized to the 15-s baseline (dF/F0). To parse electrodes into active and inactive populations, we used a threshold for GCaMP calcium activity of +3 standard deviations over the baseline measured activity. Cells that exceeded this baseline were considered “active.” Active cells were identified as “continuous” if they exceeded threshold for at least 0.5 s continuously, and “non-consecutive” cells if not. Throughout the text, we refer to “continuous active” neurons simply as “active” and “non-consecutive active” cells as “non-consecutive.”

### Separation of excitatory, inhibitory, and uncertain populations

We found that neurons could not be easily separated into populations that expressed red fluorescence (inhibitory) or did not (excitatory) based on a simple threshold because the red intensity in all neurons was continuous and not bimodal (SFig. 1A). We found that the baseline in both the red and the green channel had a strong linear relationship, implying that the intensity in the green channel influenced the intensity in the red channel. We confirmed the fluorescent contamination by measuring intensity in the red channel during ICMS, in which a change in “red intensity” was detectable during ICMS, indicating an influence of activity-dependent GCaMP on the measured red intensity (SFig. 1B). While fluorescent contamination could be due to fluorescence resonance energy transfer, more likely it is related to overlap in the emissions spectra in GCaMP and tdTomato and the applied red and green filters. The green filter applied is an et525 filter which bandpasses 500-550 nm. The red filter applied is an et595 filter with a band pass range of 570-620 nm. GCaMP has maximum emissions at 514 nm, but still has 10% max emissions at 570 nm and 1% max emissions at 620 nm, which means that GCaMP emissions can influence red channel detection. tdTomato has max emissions at 580 nm and has 10% of max emissions at 550 nm which drops to 0% at 530 nm. The overlap in the tdTomato and the green filter then was minimal and we do not expect the red channel to have strong influence on detected green fluorescence. However, all green intensity data was normalized to baseline, so any influence of the red channel on the green channel should be accounted for in this analysis.

To account for the apparent influence of activity in the green channel on red fluorescence, we used linear regression to create a model describing the relationship between baseline in the green channel and baseline in the red channel (SFig. 1C). Given that we expect there to be less inhibitory neurons present (because inhibitory neurons make up a smaller percent of cortical neurons) and because we expect neurons on the fit line to have red intensity strongly related to baseline green intensity, we only selected neurons that were greater than the root mean squared error above the fit line as “inhibitory neurons.” All neurons below the fit line were considered to be “excitatory neurons.” Neurons in between the line and the line plus the root mean squared error were classified as “uncertain” because we expect some of these neurons to be excitatory and some to be inhibitory. Indeed, we see that the “uncertain” population averages often fall in between the inhibitory and excitatory populations with a bias towards the excitatory average. We compared this simple separation method to other more complex methods that tried to optimize separation by adjusting the slope or intercept of the model (SFig. 1D,E). We found that, while all methods led to similar trends, the simple regression method yielded the strongest (or similar) separation in the average traces, so we applied this method throughout the paper. It is also worth noting that many excitatory neurons were not visible until they were activated, indicating that not all excitatory neurons were identified (while non-firing inhibitory neurons could still be identified by tdTomato).

### Classifying individual neurons responses

Neurons were divided into classes based on temporal characteristics of the calcium-evoked responses during ICMS similar to previous work^13–15^. Criteria for separation were selected based on principles derived from visual observation of individual traces to result in reasonable separation of the different observed profiles. Neurons were divided based on if they increased across the ICMS train (facilitating), remained stable across the ICMS train (non-adapting), adapted across the ICMS train (adapting), or decreased below baseline during ICMS (decrease). Adapted neurons were further divided based on the rate of adaptation, into “rapidly adapting” and “slowly adapting,” similar to previous work^13^. To meaningfully separate these classes, we compared the change in activation over time to the noise in baseline signal. “Facilitating” neurons were defined as neurons with a max peak at the end of ICMS (30-33 s) that was greater than the max peak at the beginning of ICMS (0-3 s) by three times the standard deviation of the baseline. “Adapting” neurons were defined as neurons that had a max peak at the end of ICMS that was less than the max peak at the beginning of ICMS by half the standard deviation. “Rapidly-adapting” (RA) neurons had <50% of their max peak after only 10 s of stimulation, while “Slowly-adapting” (SA) neurons had >50% of their max peak at 10 s of stimulation. “Decreased” neurons were defined as having mean activation across the entire response that was less than the baseline average by half the standard deviation. “Non-adapting” (NA) neurons were neurons that were not classified as adapting, facilitating, or decreased.

In addition to classifying neurons based on the response during ICMS, we also classified neurons based on their response after ICMS. We observed from individual traces that neurons generally fell into four categories of responses: a) activity that was below baseline after ICMS (post-ICMS depression), b) activity that returned to baseline after ICMS (post-ICMS baseline), c) activity that was elevated during the entire post-ICMS period (post-ICMS elevation), or d) rebound excitation after ICMS (post-ICMS rebound). For post-ICMS depression, mean activity during the post-ICMS period was less than the negative mean during the baseline minus half the standard deviation. For post-ICMS elevation, mean activity during the post-ICMS period was greater than the positive mean during the baseline plus half the standard deviation. “Post-ICMS rebound” was a less common event in which the period immediately following ICMS (30-33 s) was near baseline but the max activity during the post-ICMS period was greater than this value by two times the standard deviation of the baseline. “Post-ICMS baseline” was defined when the activity didn’t fall into any of these categories i.e., the activity was less than half the standard deviation above or below the baseline activity. For the plots, we only included classes that made up at least 2% of neurons (since classes with less than this only represented 1-7 neurons, resulting in noisy relationships).

### Adaptation and depression indices

We were interested in understanding how activity of individual excitatory and inhibitory neurons changes both during and after ICMS. As described in the previous methods section, adapted neurons had a smaller intensity value towards the end of the train, and facilitated neurons had stronger activity at the end of the train. To quantify the magnitude of adaptation/facilitation, we used a simple equation we defined as the “adaptation index”:

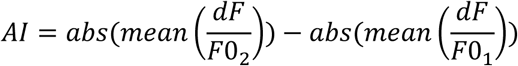

Where dF/F0_2_ represents the second half of the evoked activity during ICMS (15-30 s), and dF/F0_1_ represents the first half of the evoked activity (0-15 s).

For post-ICMS depression, we were interested in knowing how much activity changed in the short period after ICMS compared to the activation during the ICMS train. To quantify the magnitude of post-ICMS depression/elevation, we used a simple equation defined as the “depression index”:

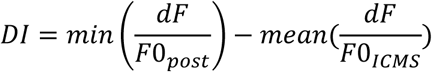

Where dF/F0_post_ represents the evoked activity in the post-ICMS period (30-50 s), and dF/F0_ICMS_ represents the evoked activity during the ICMS period (0-30 s).

### Correlation indices

To measure the synaptic connectivity of individual neurons both during (ICMS) and before (baseline, BL) ICMS, we used pairwise correlations between calcium traces for individual neurons using the *corrcoef* function in MATLAB. Traces were normalized so that correlation values represent similarities in the temporal shape of the calcium profile rather than the magnitude. Individual neurons were selected within specified distance bins away from the electrode (<200 or >200 µm). Each individual neuron then had correlation indices calculated for all other neurons of a specific cell type (excitatory or inhibitory) and the distances between the base neuron and the comparison neurons were calculated, and then we plotted the average and standard error within each of these distance bins (0-25, 25-50, 50-100, 100-200, 200+ µm). To ensure that correlation metrics were measuring real correlations in neural activity, we also performed correlations of individual neurons to white noise (WN) produced using the *wgn* function in MATLAB that was processed using the same filters (described in *Image Processing*) as the neural data. For neuron-neuropil correlations, we similarly calculated pairwise correlations of the individual neuron trace to the activity in an annulus surrounding the ROI. The diameter of the inner ring of the annulus was 2x the diameter of the ROI (leaving some space between the ROI and the annulus) and the diameter of the outer ring was 3x the diameter of the ROI.

### Statistics

All statistical analyses were conducted in MATLAB (Mathworks, Natick, MA). Unless otherwise noted, all groups were compared using ANOVA with Tukey’s HSD for significant differences (accounting for multiple comparisons). Values are presented at throughout the text as mean±standard error. We considered values to be significant at α = 0.05. Throughout the figures, *p* < 0.05 = *, *p* < 0.01 = **, *p* < 0.001 = ***.

---

**SFig. 1:**
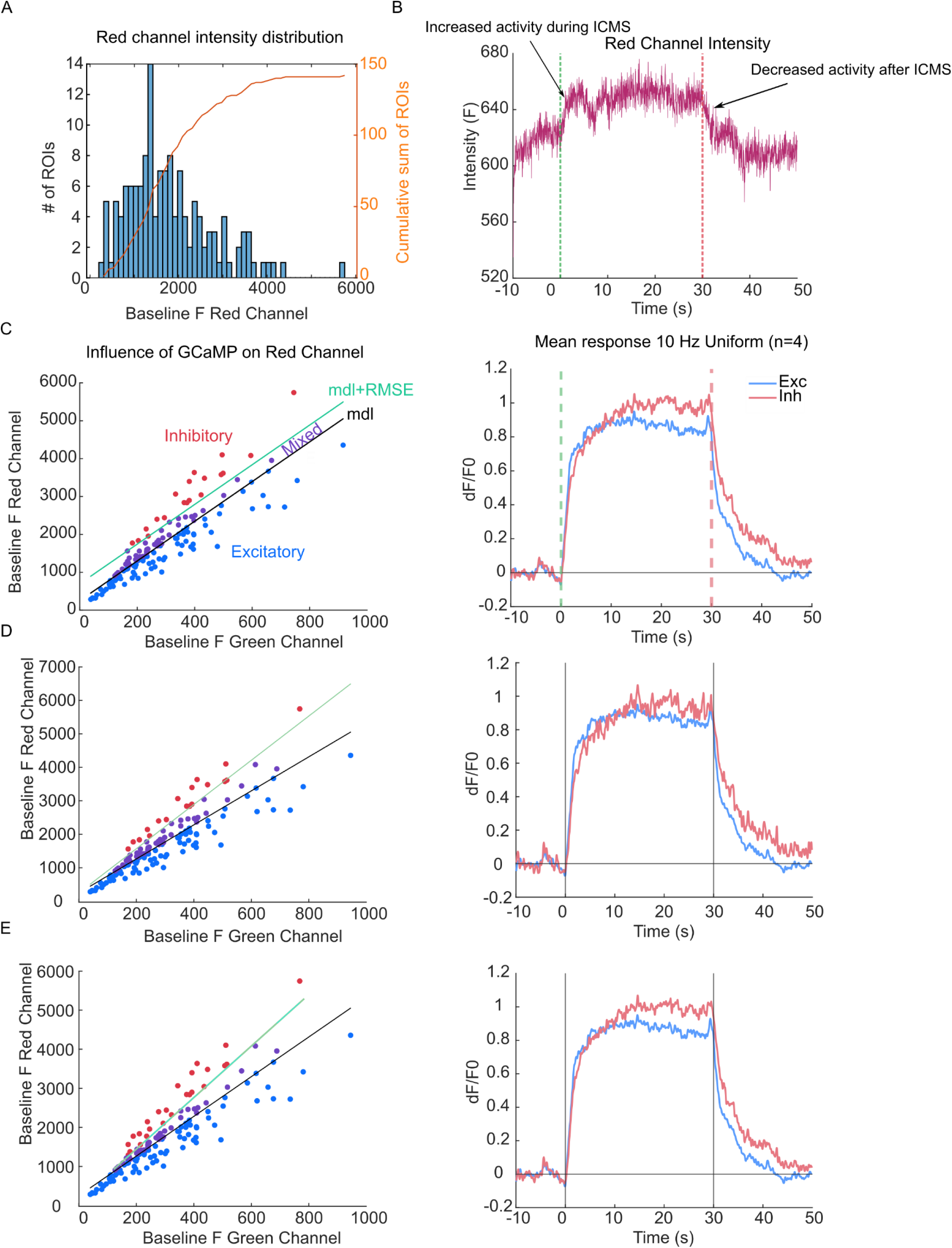
Separation of inhibitory and excitatory populations based on red channel intensity. A) Neurons did not clearly divide into a bimodal distribution (inhibitory and excitatory neurons) based on red channel intensity. The histogram shows the distribution of red fluorescence in a single animal, and the orange line shows the cumulative sum. B) We detected green channel activity in the red channel during ICMS evoked activity, indicating leakage of the green channel into the red channel. C) To account for contamination of green into red, we used a regression model, where neurons above the line plus the root mean squared error were considered inhibitory, while neurons below the regression line were considered excitatory. This separation resulted in distinct responses to ICMS among the inhibitory and excitatory subpopulations (right) D-E) We separated inhibitory and excitatory populations with slightly more complicated methods that adjusted the slope to capture an estimated 20% of inhibitory neurons (D) or calculated regression a second time after dropping points below the first regression line (E).

**SFig. 2:**
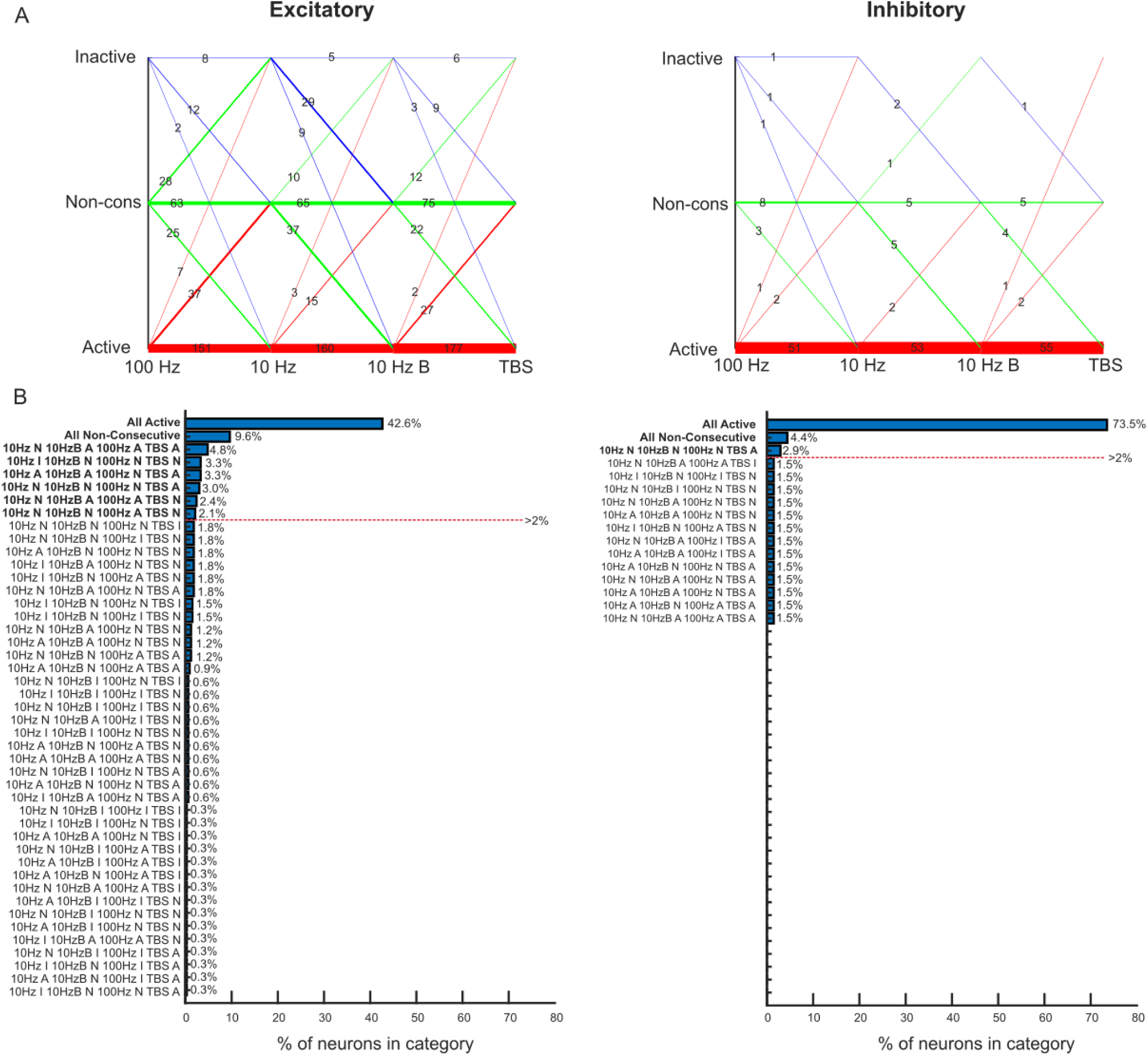
Switching of neurons between active, non-consecutively active, and inactive between the four ICMS trains. A) Each line represents neurons that switched from one category (inactive, non-cons, and active) to another between trains. The thickness and the number on the line indicates how many neurons switched between the given categories. Lines were colored based on the category they fell into for 100 Hz. B) Line plots showing the percent of neurons that fall into each possible combination of groups. Groups with bolded names indicate groups that make up more than 2% of the total number of neurons.

**SFig. 3:**
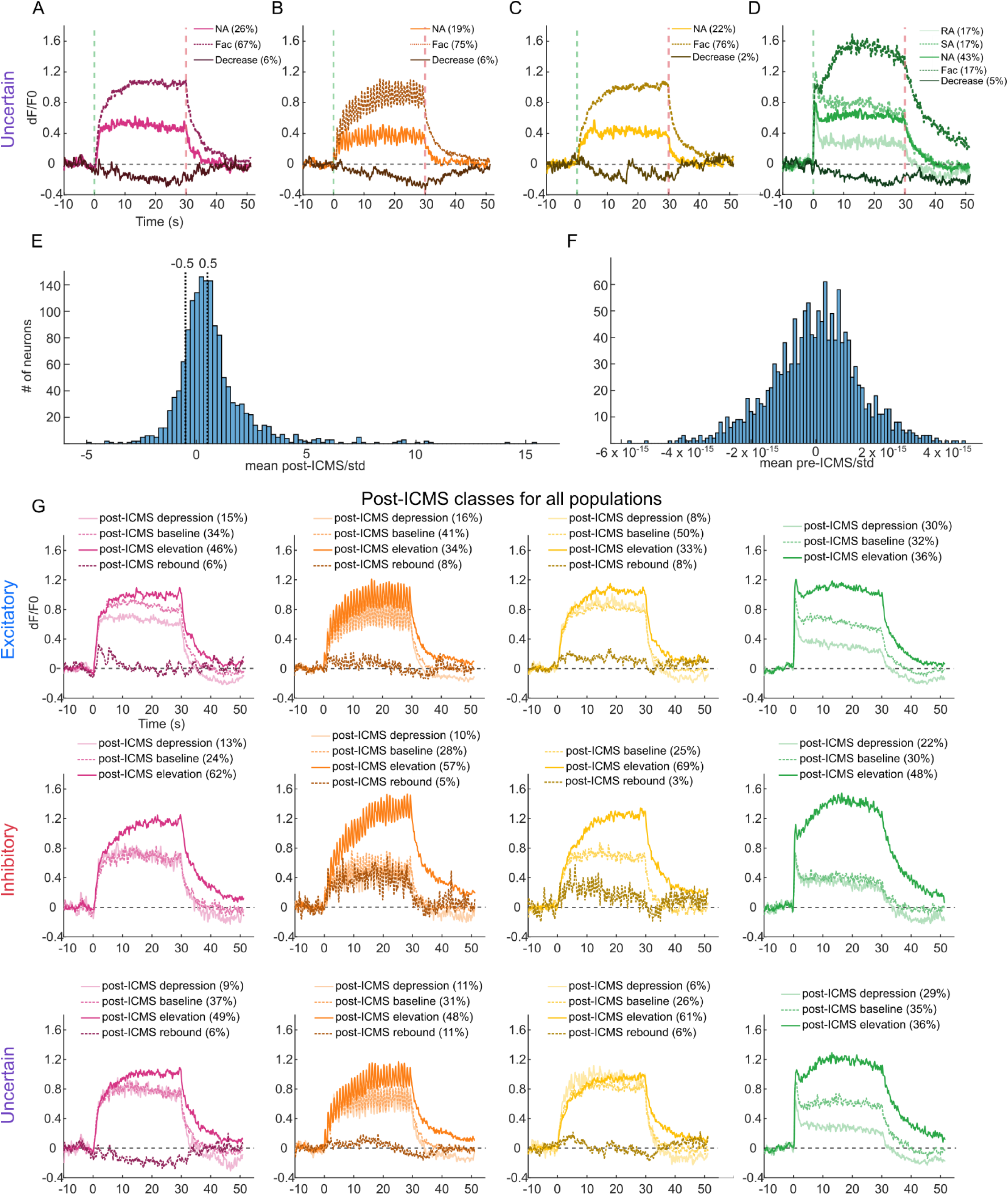
Average traces for neurons with different ICMS and post-ICMS response profiles. A-D) ICMS-evoked classes for uncertain neurons. E-F) Histograms showing the ratio of the mean activity to the standard deviation for post-ICMS (E) and pre-ICMS (F) responses. Vertical dotted lines mark -0.5 and 0.5 standard deviations away from the mean (which was used to divide the post-ICMS responses). Note that the pre-ICMS mean/baseline (F) has much smaller values than the post-ICMS mean (E) due to the large differences in activity in these two intervals. G) Individual traces for post-ICMS classes for each of the four trains. The legend indicates the percent of neurons in each class.

**SFig. 4:**
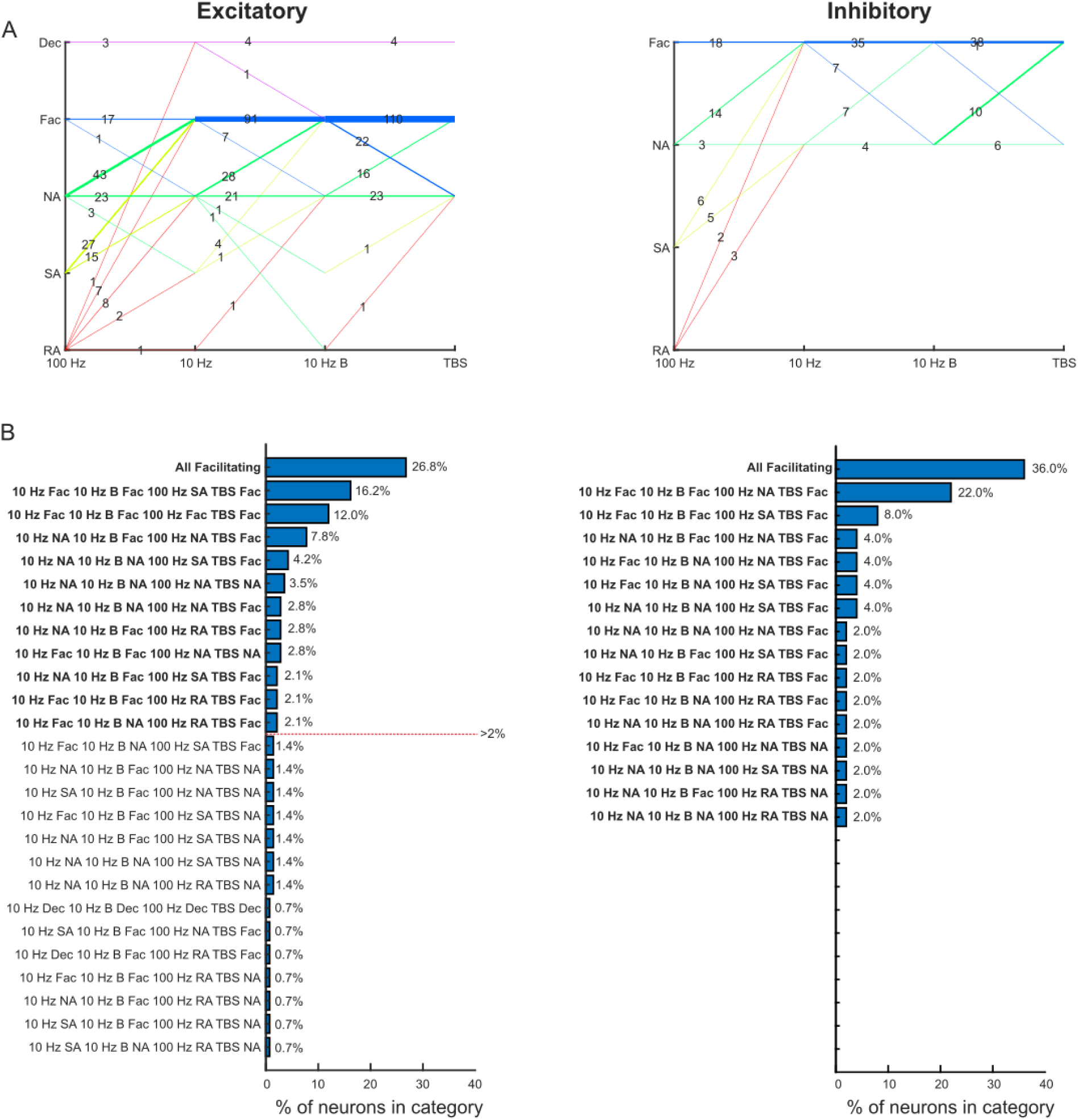
Switching of neurons between individual classifications (RA, SA, NA, Fac, or Dec) between the four ICMS trains. A) Each line represents neurons that switched from one category (RA, SA, NA, Fac, or Dec) to another between trains. The thickness and the number on the line indicates how many neurons switched between the given categories. Lines were colored based on the category they fell into for 100 Hz. B) Line plots showing the percent of neurons that fall into each possible combination of groups. Groups with bolded names indicate groups that make up more than 2% of the total number of neurons.

**SFig. 5:**
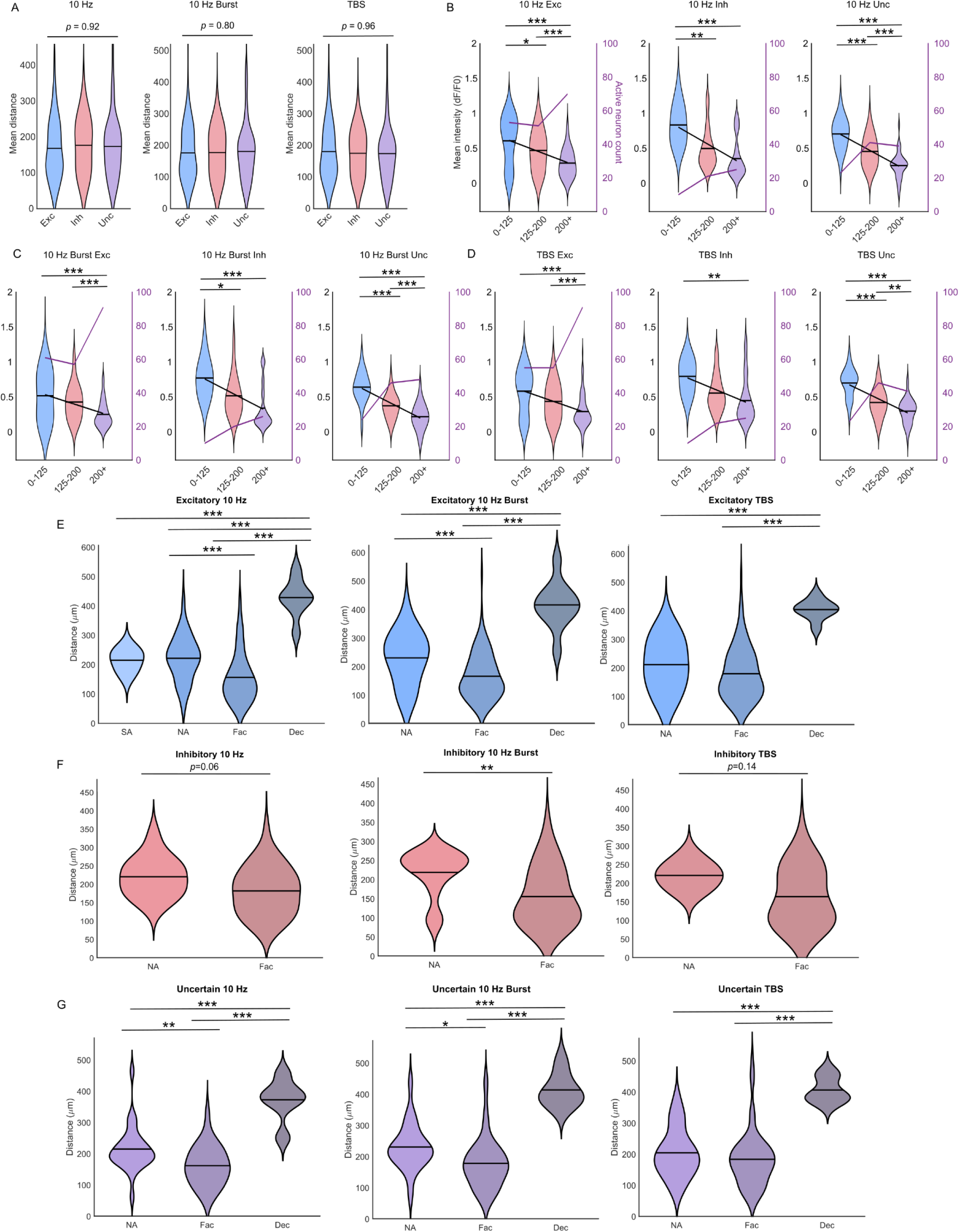
Relationships between subpopulation responses and distance from the electrode. A) There were no significant differences for the distance of each subpopulation from the electrode. B-D) Mean intensities for neurons within different distance bins in response to 10 Hz (B), 10-Hz Burst (C), and TBS (D). For all subpopulations and trains, the mean intensity decreased farther from the electrode. E-G) Mean distances from the electrode for different classes of neurons based on temporal response for excitatory (E), inhibitory (F), and uncertain (G) populations.

**SFig. 6:**
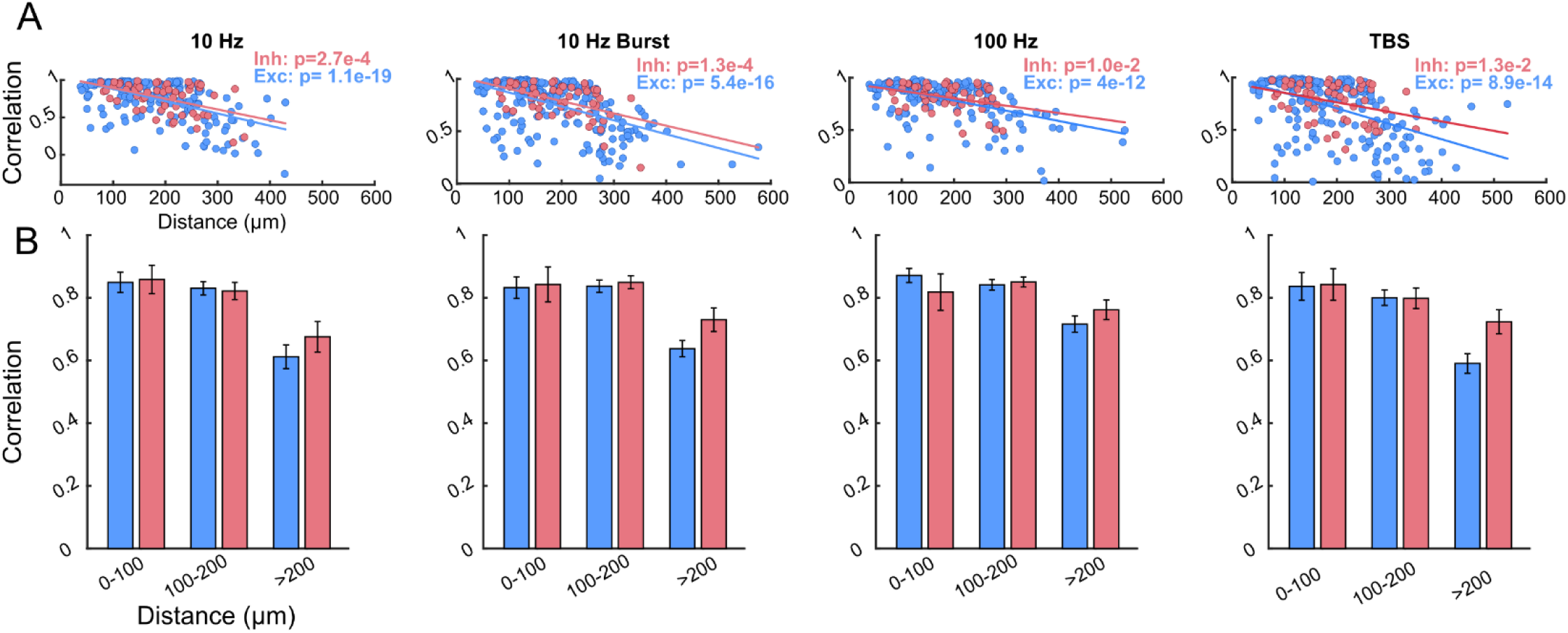
Plots for neuron-neuropil correlation for individual ICMS trains. A) Individual neuron correlations plotted as a function of distance from the electrode. The solid lines represent linear fits to the data. B) Average neuron to neuropil correlations in different distance bins. Error bars represent the standard error.

## Notes

https://www.bioniclab.org

### Competing Interest Statement

The authors have declared no competing interest.

